# *Tsc1* Deletion in Purkinje Neurons Disrupts the Axon Initial Segment, Impairing Excitability and Cerebellar Function

**DOI:** 10.1101/2025.01.31.635932

**Authors:** Samuel P. Brown, Achintya K. Jena, Joanna J. Osko, Joseph L. Ransdell

**Author notes:** Correspondence, Miami University, 700 E High St. Oxford, OH 45056. Authors contributed equally (Samuel P. Brown, Achintya K. Jena).

## Abstract

Loss-of-function mutations in tuberous sclerosis 1 (*TSC1*) are prevalent monogenic causes of autism spectrum disorder (ASD). Selective deletion of *Tsc1* from mouse cerebellar Purkinje neurons has been shown to cause several ASD-linked behavioral impairments, which are linked to reduced Purkinje neuron repetitive firing rates. We used electrophysiology methods to investigate why Purkinje neuron-specific *Tsc1* deletion (*Tsc1^mut/mut^*) impairs Purkinje neuron firing. These studies revealed a depolarized shift in action potential threshold voltage, an effect that we link to reduced expression of the fast-transient voltage-gated sodium (Nav) current in *Tsc1^mut/mut^* Purkinje neurons. The reduced Nav currents in these cells was associated with diminished secondary immunofluorescence from anti-pan Nav channel labeling at Purkinje neuron axon initial segments (AIS). Interestingly, anti-ankyrinG immunofluorescence was also found to be significantly reduced at the AIS of *Tsc1^mut/mut^* Purkinje neurons, suggesting Tsc1 is necessary for the organization and functioning of the Purkinje neuron AIS. An analysis of the 1^st^ and 2^nd^ derivative of the action potential voltage-waveform supported this hypothesis, revealing spike initiation and propagation from the AIS of *Tsc1^mut/mut^* Purkinje neurons is impaired compared to age-matched control Purkinje neurons. Heterozygous *Tsc1* deletion resulted in no significant changes in the firing properties of adult Purkinje neurons, and slight reductions in anti-pan Nav and anti-ankyrinG labeling at the Purkinje neuron AIS, revealing deficits in Purkinje neuron firing due to *Tsc1* haploinsufficiency are delayed compared to age-matched *Tsc1^mut/mut^* Purkinje neurons. Together, these data reveal the loss of *Tsc1* impairs Purkinje neuron firing and membrane excitability through the dysregulation of proteins necessary for AIS organization and function.

## INTRODUCTION

Tuberous sclerosis complex (TSC) is an autosomal dominant disorder affecting multiple organ systems, which causes severe and progressive deficits in nervous system functioning (Crino et al., 2006), often resulting in seizures (Holmes et al., 2007), cognitive deficits (Marcotte and Crino, 2006), and impaired behaviors. TSC is caused by loss of function mutations in either *TSC1* or *TSC2* (European Chromosome 16 Tuberous Sclerosis Consortium, 1993; Au et al., 2007), which results in exaggerated mammalian target of the rapamycin complex 1 (mTORC1) signaling (Tee et al., 2002, 2003; Wullschleger et al., 2006). This increase in mTORC1 activity drives excessive protein synthesis through S6K1 and 4E-BP1 effector molecules and lipid synthesis through the activation of transcription factors (Fingar et al., 2002). Interestingly, 41-69% of individuals diagnosed with TSC exhibit autism-like cluster manifestations and 40-50% are diagnosed with autism spectrum disorder (ASD) (Wiznitzer, 2004; Jeste et al., 2008; de Vries et al., 2023). ASD is a neurodevelopmental disorder in which the etiology is often unknown, but can include a combination of genetic and environmental factors (Díaz-Anzaldúa and Díaz-Martínez, 2013; Port et al., 2014). ASD impairments are strongly linked with deficits in the functioning of the cerebellum (Wang et al., 2014; Stoodley et al., 2017; Gibson et al., 2023). Post-mortem studies of individuals diagnosed with ASD have linked the disorder with reduced numbers of cerebellar Purkinje neurons (Bailey et al., 1998; Stoodley, 2014) and in animal models, attenuated excitability in cerebellar Purkinje neurons has been directly linked to ASD-like behavioral phenotypes (Kalume et al., 2007; Tsai et al., 2012; Cupolillo et al., 2016; Peter et al., 2016). Across cerebellar circuits, Purkinje neurons function to integrate incoming sensory information and are the sole output neurons of the cerebellar cortex (Palkovits et al., 1972; Andersen et al., 1992). Purkinje neurons fire repetitive action potentials spontaneously, providing consistent GABAergic inhibition to deep cerebellar nuclei neurons (Ito et al., 1964; Obata et al., 1967, 1970). Studies using animal models have consistently revealed targeted disruptions to Purkinje neuron intrinsic firing properties drive a multitude of impairments in behaviors reliant on cerebellar functioning (Levin et al., 2006; Tsai et al., 2012, 2018; Bosch et al., 2015; Peter et al., 2016; Ransdell et al., 2017).

To investigate the role of Tsc1 in the functioning of cerebellar Purkinje neurons, a Cre-recombinase-*LoxP* recombination strategy (Orban et al., 1992) was used to selectively delete *Tsc1* from mouse Purkinje neurons (Barski et al., 2000; Zhang et al., 2004; Tsai et al., 2012). Interestingly, the homozygous deletion of *Tsc1* in Purkinje neurons resulted in mice, referred to here as *Tsc1^mut/mut^*, with several ASD-linked behavioral phenotypes that included impairments in motor functioning, social interactions, vocalizations, and exaggerated repetitive behaviors. These behavioral deficits were linked to attenuated repetitive firing in *Tsc1^mut/mut^* Purkinje neurons and eventual (and progressive) Purkinje neuron apoptosis (Tsai et al., 2012, 2018; Lawson et al., 2024). Heterozygous *Tsc1* deletion from Purkinje neurons (*Tsc1^mut/+^*) was also shown to cause attenuated Purkinje neuron firing and ASD-like behavioral phenotypes (Tsai et al., 2012, 2018; Lawson et al., 2024), although these impairments are less severe than those measured in *Tsc1^mut/mut^* mice. Attenuated intrinsic excitability of *Tsc1^mut/mut^* Purkinje neurons could be rescued via intraperitoneal injections of rapamycin, an acute inhibitor of mTORC1 signaling, if rapamycin treatments were started by 6 weeks of age, suggesting the impaired firing of *Tsc1^mut/mut^* Purkinje cells may reflect changes in the expression and/or gating properties of ion channels (Tsai et al., 2018).

We investigated the underlying causes of the reduced repetitive firing in *Tsc1^mut/mut^* Purkinje neurons. While *Tsc1* deletion is often associated with exaggerated protein expression and cell growth (Wullschleger et al., 2006), our experiments reveal *Tsc1^mut/mut^* Purkinje neurons have diminished anti-pan Nav channel immunofluorescence at the Purkinje neuron AIS, which corresponds with reduced Nav currents and impaired action potential initiation and propagation at the axon initial segment (AIS). At the AIS of *Tsc1^mut/mut^* Purkinje neurons, we also measured reduced immunofluorescence signals from anti-ankyrinG labeling. AnkyrinG is a critical cytoskeletal regulator of AIS segment organization and function, which has been directly linked to the recruitment and clustering of Nav channels at the AIS of Purkinje neurons (Zhou et al., 1998; Jenkins and Bennett, 2001). These data shed light on the pathophysiology of the *Tsc1^mut/mu^*^t^ mouse model and provide new insights into how loss-of-function mutations in *TSC1* may lead to deficits in neuronal membrane excitability and circuit function.

## Methods

### Animals

All animal experiments were performed in accordance with protocols approved by the Miami University Institutional Animal Care and Use Committee guidelines. Experiments utilized male and female wild type C57BL/6J mice and transgenic lines with a C57BL/6J strain background. For neonatal dissociation experiments, animals were P15-16 and sex was unknown. For all other experiments (slice current-clamp, slice voltage-clamp, and immunofluorescence), animal numbers, sex, and ages are described in Supplemental Table 1. Within each genotype (wild type, *Tsc1^mut/+^*, and *Tsc1^mut/mut^*), no sex-specific differences were measured in firing frequency (Supplemental Figure 1) or other action potential properties. Sex-specific differences were not assessed in voltage-clamp and immunofluorescence experiments due to insufficient animal numbers (see Supplemental Table 1).

*Tsc1* was selectively deleted from mouse cerebellar Purkinje neurons by crossing *Tsc1* floxed animals (*Tsc1^flox/flox^*, Kwiatkowski et al., 2002) (Jackson laboratory, strain # 005680) with hemizygous transgenic animals expressing Cre-recombinase on the L7/Pcp2 promoter-strain B6.Cg-Tg(Pcp2-cre)3555Jdhu/J (Zhang et al., 2004) (Jackson laboratory, strain # 010536). Cre-recombinase positive animals were also crossed with the Ai14 Cre-reporter strain (Jackson Laboratory, strain # 007914), resulting in tdTomato expression in Cre-recombinase positive cells (see Supplemental Figure 2).

### Control Groups

Age-matched wild type animals were used as controls for current-clamp electrophysiology experiments. Purkinje neurons from L7/Pcp2 Cre-positive animals, as well as *Tsc1^flox/flox^*;Cre-negative animals, were previously shown to have similar repetitive firing properties as wild type Purkinje neurons (Tsai et al., 2012). For voltage-clamp experiments using dissociated neonatal cells, control cells were taken from L7/Pcp2 Cre-positive animals. In voltage-clamp experiments using adult Purkinje neurons (in acute cerebellar slices), control group cells were taken from both wild type animals as well as *Tsc1^flox/flox^*; Cre-negative animals. Nav current properties in these control groups were determined to be statistically similar (Supplemental Figure 3) and reported results from these control groups are combined. Immunofluorescence studies used both wild type and L7/Pcp2 Cre-positive;Ai14 Cre-reporter animals as control groups. Immunofluorescence intensity measures associated with anti-pan Nav and anti-ankyrinG labeling were also determined to be similar between control groups (Supplemental Figure 4) and data from these control groups are combined.

### Preparation of acute cerebellar slices

Mice were anesthetized using an intraperitoneal injection of 1 mL/kg ketamine (10 mg/mL)/xylazine (0.25 mg/mL) cocktail and perfused transcardially with 25 mL cutting solution containing (in mM): 240 sucrose, 2.5 KCl, 1.25 NaH_2_PO_4_, 25 NaHCO_3_, 0.5 CaCl_2_, and 7 MgCl_2_. Brains were rapidly dissected, glued to a specimen tube, and submerged in warmed agarose dissolved in cutting solution. 350 µm parasagittal slices were cut in ice-cold cutting solution saturated with 95% O_2_/5% CO_2_ using a Compresstome VF-300 vibratome (Precisionary Instruments). Slices were placed on stretched nylon mesh in oxygenated artificial cerebrospinal fluid (ACSF) containing (in mM): 125 NaCl, 2.5 KCl, 1.25 NaH_2_PO_4_, 25 NaHCO_3_, 2 CaCl_2_, 1 MgCl_2_, and 25 dextrose at pH 7.4 (∼300 mOsM/L) for 25 minutes at 33 °C. Slices were then incubated in room temperature ACSF for at least 35 minutes before electrophysiological recordings. Purkinje neurons were identified in the Purkinje neuron layer of cerebellar sections using a SliceScope Pro 3000 (Scientifica). During electrophysiology experiments, warmed (32-33 °C) oxygenated-ACSF was continuously perfused.

### Acute dissociation of neonatal Purkinje neurons

For neonatal voltage-clamp experiments, mice were anesthetized using an intraperitoneal injection of a ketamine/xylazine cocktail as described above. Post-anesthesia, mice were rapidly decapitated, and the brain was rapidly dissected and placed in ice-cold dissociation solution containing (in mM): 82 Na_2_SO_4_, 30 K_2_SO_4_, 5 MgCl_2_, 10 HEPES, 10 glucose, and 0.001% phenol red (at pH 7.4). A scalpel was used separate and mince the cerebellar vermis into small tissue chunks. Cerebellar tissue was then transferred to dissociation solution containing 3 mg/ml protease XXIV at 33 °C for 15 minutes. Minced cerebellar tissue was then transferred to dissociation solution containing 1 mg/mL bovine serum albumin and 1 mg/mL trypsin inhibitor and incubated at room temperature for 5 minutes before being transferring to ACSF saturated with 95% O_2_/5% CO_2_ on stretched nylon mesh for 35 minutes. Before recordings, an aliquot of cerebellar tissue was dissociated by triturating tissue pieces with fire-polished glass pipettes.

### Current- and voltage-clamp recordings

Whole-cell patch-clamp recordings used glass microelectrodes (borosilicate standard wall, 1.5 mm outer diameter, 0.86 mm inner diameter, Harvard Bioscience) with 2-4 MΩ resistance values, pulled on the day of the experiment using a P-1000 Flaming/Brown Micropipette Puller (Sutter Instrument). Electrodes for current-clamp studies were filled with an internal solution containing (in mM): 144 K-gluconate, 0.2 EGTA, 3 MgCl_2_, 10 HEPES, 4 MgATP, 0.5 NaGTP, at pH 7.4 (∼300 mOsM/L). Prior to patching, electrode tip potentials were zeroed. During current-clamp experiments, recorded voltages were corrected for a 17.5 mV liquid junction potential in real-time. Spontaneous and evoked action potentials were recorded using a dPatch amplifier and SutterPatch software (Sutter Instruments). Action potential threshold voltages were calculated as the voltage measured when dV/dt is >10 mV/100 µs or when 25% of the maximum action potential dV/dt is reached, whichever is smaller. Action potential duration (APD) was calculated as the time interval from the pass of the threshold voltage during the action potential upstroke until the pass of threshold voltage during action potential downstroke. Each action potential measurement from a given cell was recorded as the average across 20 consecutive action potentials in a spontaneously firing cell. Evoked action potentials were recorded during a current-clamp protocol with an initial −500 pA (100 ms) current injection before steps to 0 pA, or other depolarizing current steps, for 700 ms. To measure the delay between spike generation in the AIS (occurring first) and secondary spike initiation in the somatic compartment, the second derivative of the voltage signal (dV^2^/dt) was plotted against time using Clampfit software (Molecular Devices). In this second derivative plot, two positive peaks, reflecting spike initiation at the AIS and somatic compartment, respectively, occur in time with the action potential upstroke. Difference in time between these peaks (delay) was measured and used for analysis.

Whole-cell voltage-clamp recordings in adult slice Purkinje neurons and in dissociated neonatal Purkinje neurons were performed at room temperature (22 °C) and used 1-3 MΩ microelectrodes filled with internal solution containing (in mM): 105 CsCl, 15 TEA-Cl, 5 4-AP, 1 CaCl_2_, 2 MgCl_2_, 10 EGTA, 4 MgATP, 10 HEPES, and 8 NaCl at pH 7.4 (∼300 mOsM/L). In these experiments, Nav currents were recorded in a reduced sodium ASCF containing (in mM): 50 NaCl, 75 TEA-Cl, 2.5 KCl, 1.25 NaH_2_PO_4_, 25 NaHCO_3_, 2 CaCl_2_, 1 MgCl_2_, 0.2 CdCl_2_, and 25 dextrose (∼300 mOsM/L), saturated with 95% O_2_/5% CO_2_. In recordings from adult Purkinje neurons in acute cerebellar slices, Nav currents were measured under appropriate space-clamp conditions using a prepulse protocol that was developed by Milescu et al. (2010) and previously used to measure Nav currents in adult Purkinje neurons in cerebellar slices (Bosch et al., 2015; Ransdell et al., 2017). To measure the voltage-dependence of Nav current activation, cells were first depolarized (typically to 0 mV) for 1 ms to drive activation and inactivation of Nav channels in the somatic compartment as well as in the distal neurites. A subsequent repolarizing step to −60 mV (for 1 ms) enabled recovery of a portion of these inactivated Nav channels, and a final depolarizing step was used to measure the Nav current properties of recovered Nav channels (at varying voltages) under appropriate space-clamp conditions. To measure the voltage-dependence of Nav current steady-state inactivation, a similar prepulse strategy was used, however, in this protocol, the repolarizing step driving Nav channel recovery from inactivation was varied and used as a conditioning voltage step. Evoked Nav currents were measured after the conditioning voltage step during a final depolarizing step to −20 mV. The persistent/steady-state component of the evoked Nav currents were digitally subtracted prior to analysis of the fast-transient Nav current (I_NaT_). I_NaT_ inactivation kinetics was assessed by fitting the inactivating portion of I_NaT_ with a first-order exponential function. From these fits, the time constant of current decay (tau, τ) was recorded across voltages. Normalized Nav current (I/I_Max_) and normalized Nav conductance (G/G_Max_) are plotted as a function of the conditioning voltage or the test voltages, respectively, to assess the voltage-dependence of Nav channel activation and steady-state inactivation. These plots were fitted using the Boltzmann sigmoidal equation below:

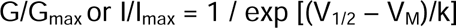

where V_M_ is the membrane potential, V_1/2_ is the membrane potential of half-maximal activation and k is the slope factor.

### Imaging

Secondary antibody immunolabeling was used to measure the fluorescence intensity of anti-pan Nav and anti-ankyrinG labeling at the AIS of cerebellar Purkinje neurons. Adult mice were anesthetized with 1 mL/kg intraperitoneal injection of ketamine/xylazine cocktail and perfused transcardially with 1X PBS followed by 1 % formaldehyde in 0.1 M PB at pH 7.4. After dissection, the cerebellum was separated from the cortex and was embedded in optimal cutting temperature compound (OCT, Fisher HealthCare) and then frozen. Sagittal cerebellar sections (25 µm) were dry mounted onto positively-charged glass slides (Epredia) using a Leica cryostat at −14 to −16 °C. Wells around tissue sections were created using clear nail polish in preparation for antibody labeling.

Cerebellar sections were washed with 1X PBS three times and then incubated in a blocking buffer containing 7.5 % goat serum and 0.25 % Triton X-100 (Fisher BioReagents) for 1-1.5 hours at 4 °C. After decanting, primary antibodies were incubated on tissue sections overnight (mouse anti pan Nav: 1:1000, Sigma-Aldrich; mouse anti-ankyrinG (IgG2a): 1:500, Sigma-Aldrich; rabbit anti-Calbindin: 1:20000, Novus) at 4 °C. Primary antibodies were then decanted and washed three times with 1X PBS, and then incubated with a solution that contained secondary antibodies and 1 % BSA and 0.25 % Triton X-100 for 1.5 hours at room temperature (goat anti-mouse Alexa 488: 1:250, Sigma-Aldrich; goat anti-mouse CF 647: 1:500, Sigma-Aldrich; goat anti-rabbit CF 568: 1:167, Sigma-Aldrich). After decanting the final time, tissue sections were washed with 1X PBS three times and a 5 minute incubation period after each wash. Slides were dried and mounted using Vectashield (Vector Laboratories) and a glass coverslip. Calbindin secondary immunolabeling or tdTomato expression in Cre-positive animals (from the Ai14 Cre-reporter strain) were used to identify AIS labeling of Purkinje neurons.

Confocal microscopy images were acquired using a Zeiss LSM-710 microscope utilizing a 63X oil-immersion or 20X objective lens. Z-stack images were acquired at a fixed interval of 1 µm and the images were analyzed with ImageJ (NIH) using maximum intensity projected CZI files.

Fluorescence intensity was measured by drawing line scans of secondary anti-ankyrinG labeling along the AIS, beginning at the somatic compartment and ending at the termination of fluorescence signal associated with anti-ankyrinG labeling. Background fluorescence was subtracted by measuring the mean secondary antibody fluorescence in the somatic compartment and subtracting this value from each sampled intensity value along the AIS. Fluorescence intensity was plotted against distance (from the beginning of the line scan). For each cell, a 5-point rolling mean was used to determine final intensity values (after background subtraction). To calculate mean intensity values, measurements of distance were aligned across cells at the sample corresponding to 10% of the peak anti-ankyrinG fluorescence signal (nearest the soma). Area under the fluorescence intensity curve (AUC) was calculated for each cell using Prism (GraphPad) and compared across experimental groups.

### Statistical analysis

Electrophysiological data and action potential analyses were performed using SutterPatch software. Current traces were analyzed using Clampfit in the pCLAMP 11 software suite (Molecular Devices). Statistical analyses were performed using Prism (GraphPad). Tests, P values, cell numbers, and animal numbers are described in Figure legends. Repeated measure (RM) two-way ANOVAs used Geisser-Greenhouse’s correction. Prior to running unpaired Student’s t-tests, an F-test was used to determine if group variance was similar. If the F-test P-value was ≤.05, the unpaired Student’s t-test included a Welch’s correction, which is noted in the text.

## RESULTS

### *Tsc1* deletion in Purkinje neurons results in impaired action potential generation

To selectively delete *Tsc1* from mouse cerebellar Purkinje neurons, a Cre-*LoxP* recombination strategy (depicted in Fig. 1A) was used. Hemizygous L7/*Pcp2*-Cre animals, which express Cre recombinase under the control of the mouse Purkinje cell-specific L7 promoter (Zhang et al., 2004; Saito et al., 2005), were crossed with *Tsc1* mutant animals with *loxP* sites flanking exons 17 and 18 of the *Tsc1* gene. To verify selective Cre expression in the Purkinje neuron layer, progeny from this cross were bred with the Ai14 tdTomato Cre-reporter strain (Jackson Laboratory, strain # 007914; Madisen et al., 2010). From these crosses, we verified robust and selective Cre-mediated tdTomato expression in neonatal (Fig. 1 *A1.*) and adult Purkinje neurons (see Supplemental Figure 2) in cerebella isolated from male and female animals.

**Figure 1.**
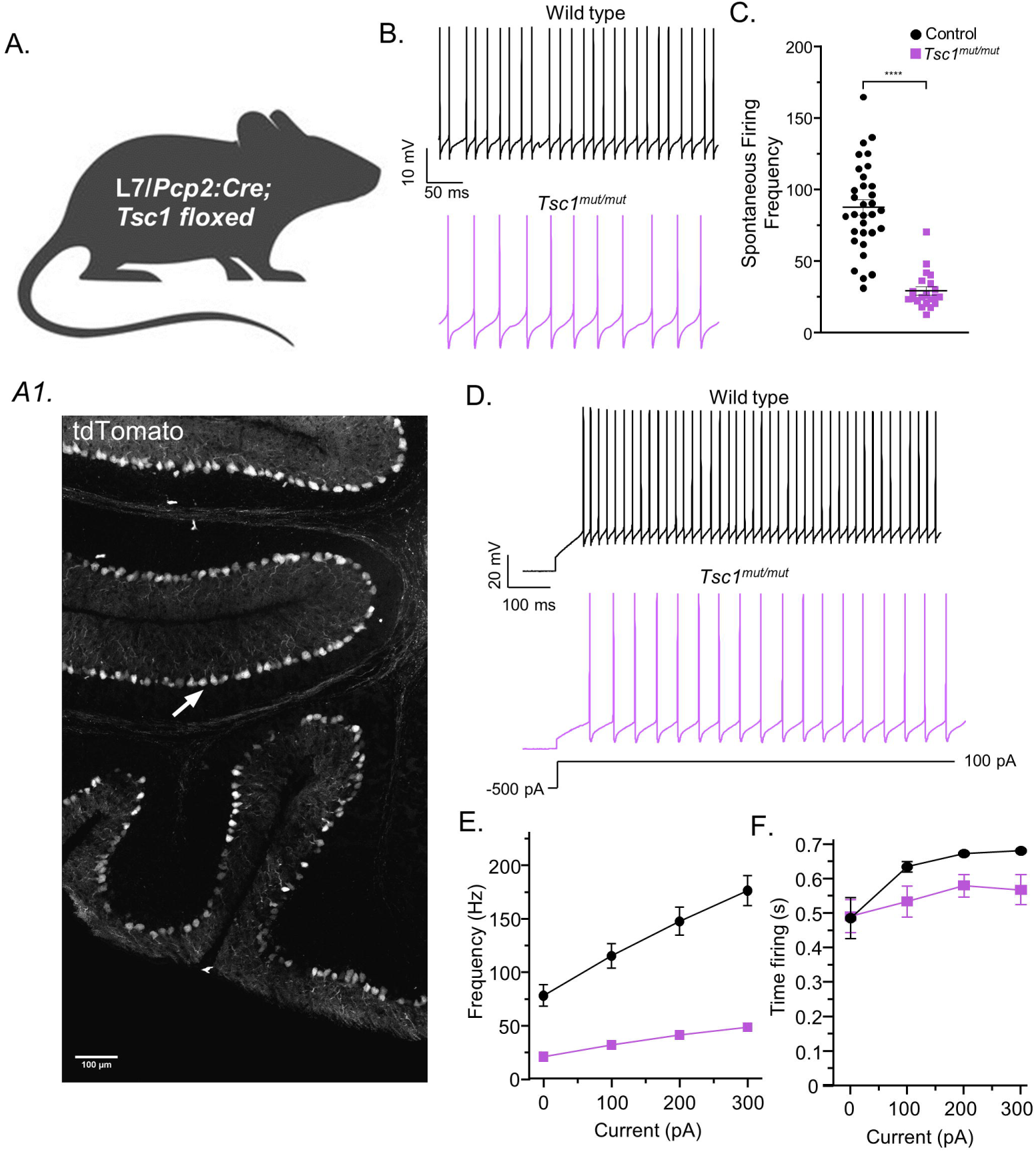
*Tsc1* deletion causes reduced repetitive firing in adult cerebellar Purkinje neurons. **A.** *Tsc1* was selectively deleted from mouse cerebellar Purkinje neurons by crossing *Tsc1* floxed mice with a transgenic line in which the L7/Pcp2 promoter directs hemizygous Cre-recombinase expression (see Methods). To verify Purkinje neuron specific Cre-recombinase expression, Cre-positive animals were also crossed with a Cre-reporter strain in which Cre-mediated recombination drives tdTomato expression. In panel *A1*, robust and selective tdTomato fluorescence is found in the Purkinje neuron layer of a sagittal cerebellar slice (see Methods) taken from a neonatal (P15) Cre-positive animal. **B.** Representative spontaneous action potential records are shown from Purkinje neurons in acute cerebellar slices from adult control (top, *black*) and *Tsc1^mut/mut^* (bottom, *magenta*) animals. **C.** In *Tsc1^mut/mut^* cells, the mean (± SEM) spontaneous firing frequency is significantly lower compared to control cells (Welch’s unpaired t-test: P < 0.0001; control N = 13, n = 32; *Tsc1^mut/mut^* N = 6, n = 21). **D.** Representative evoked firing records are also shown from control (top, *black*) and *Tsc1^mut/mut^*(bottom, *magenta*) cells. **E.** Mean (± SEM) evoked firing frequencies are plotted against current injections and reveal evoked firing in *Tsc1^mut/mut^* cells is also significantly (RM two-way ANOVA: P < .0001) attenuated compared to control cells (DF = 1, F-value = 56.1). **F.** The mean (± SEM) durations of repetitive firing during the 0.7 s depolarizing current injections are plotted against depolarizing current injection amplitude and reveals *Tsc1^mut/mut^* cells have an impaired capacity to sustain repetitive firing compared to control cells (RM two-way ANOVA: P < .05; DF = 1, F-value = 4.3).

Test group animals that were homozygous for the mutant *Tsc1* floxed allele and that expressed Cre-recombinase selectively in Purkinje neurons, referred to henceforth as *Tsc1^mut/mut^* animals, were used for patch-clamp electrophysiology studies and were compared directly with wild type controls. Parasagittal cerebellar brain sections were acutely isolated from adult (5-8 week-old) animals for whole cell Purkinje neuron current-clamp recordings at physiological temperatures. In these records, we assessed spontaneous (Fig. 1B) and evoked (Fig. 1D) Purkinje neuron firing in control and *Tsc1^mut/mut^* Purkinje neurons. These recordings revealed *Tsc1^mut/mut^* Purkinje neurons have spontaneous firing frequencies that are significantly attenuated compared to wild type Purkinje neurons (Fig. 1C). Evoked firing frequencies were measured during 0.7 second depolarizing current injections that were applied immediately after a −500 pA hyperpolarizing current injection (used to silence spontaneous electrical activity, see Fig. 1D). These protocols also revealed *Tsc1^mut/mut^* Purkinje neurons have significantly lower mean (±SEM) evoked firing frequencies when compared to the evoked firing in wild type Purkinje neurons (Fig. 1E). During evoked firing tests, the mean (±SEM) durations of repetitive firing (during each 0.7 s depolarizing current injection) were found to be significantly lower in *Tsc1^mut/mut^* Purkinje neurons, revealing a reduced capacity to maintain high rates of repetitive firing in these cells.

Using the gap-free current-clamp recordings of spontaneously firing cells, we compared action potential waveform properties, presented as representative records in Figure 2A, and identified notable differences in *Tsc1^mut/mut^* Purkinje neuron action potentials from control cells. Compared to wild type controls, the mean (±SEM) action potential threshold voltage was significantly more depolarized in *Tsc1^mut/mut^* cells (Fig. 2B). The amplitude of *Tsc1^mut/mut^* action potentials was also found to be lower than control neurons (Fig. 2C), and the mean duration of *Tsc1^mut/mut^* Purkinje neuron action potentials (APD) was significantly longer than action potentials measured in control cells (Fig. 2D). From these data, we hypothesized that changes in the expression and/or gating properties of voltage-gated ion channels were responsible for the impaired membrane excitability of *Tsc1^mut/mut^* Purkinje neurons.

**Figure 2.**
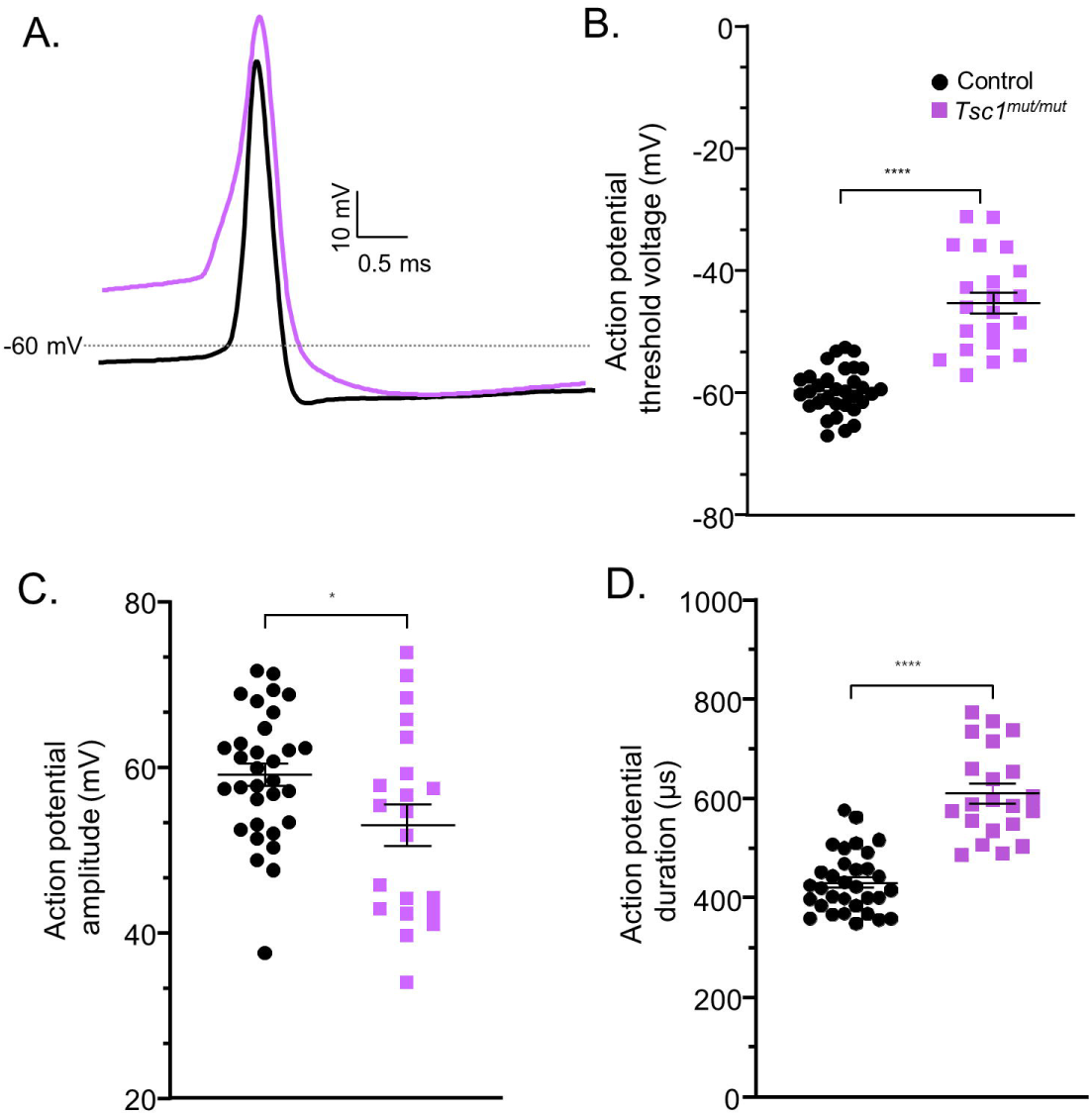
Action potential waveforms recorded from *Tsc1^mut/mut^* Purkinje neurons are significantly different from those recorded from control cells. **A.** Representative action potential records from adult control (*black*) and *Tsc1^mut/mut^* (*magenta*) Purkinje neurons are overlayed for comparison. **B.** The mean (± SEM) action potential threshold voltage is significantly more positive in *Tsc1^mut/mut^* cells (*magenta squares*) compared to control (*black circles*) cells (Welch’s unpaired t-test: P < 0.0001). **C.** The mean (± SEM) amplitude of the action potential waveform in *Tsc1^mut/mut^* cells is significantly (Welch’s unpaired t-test: p = 0.0256) reduced compared to the mean (± SEM) amplitude of control cells. **D.** Action potential duration values are significantly larger in *Tsc1^mut/mut^* cells compared to controls (Welch’s unpaired t-test: P < 0.0001), (control N = 13, n = 32; *Tsc1^mut/mut^* N = 6, n = 21).

### Loss of *Tsc1* affects Nav currents in adult cerebellar Purkinje neurons

The depolarized shift in the action potential threshold voltage suggested there may be changes in the voltage-dependence of Nav channel activation, or in the expression of voltage-gated sodium channels. To explore this hypothesis, we first measured the properties of Nav currents in Purkinje neurons acutely dissociated from neonatal (P15-16) control and *Tsc1^mut/mut^* animals. A representative micrograph of a dissociated Purkinje neuron (with patch electrode) is presented in Figure 3A. Acute dissociation of these cells enable the removal of long neurite projections, particularly the axonal projection, which improves membrane space clamp such that the very fast (and large) Nav currents can be appropriately measured. Also, to improve Nav current measurements, the extracellular ACSF (bath) included reduced (50 mM) sodium chloride and pharmacological blockers for potassium and calcium conductances (see Methods). The voltage-dependence of the fast-transient sodium current (I_NaT_) activation was measured by applying depolarizing voltage steps from a −90 mV holding potential (Fig. 3B). Measurements of the mean peak I_NaT_ across depolarizing voltage steps revealed control and *Tsc1^mut/mut^* Purkinje neurons have I_NaT_ currents which activate at similar voltages and that are similar in amplitude (Fig. 3C). I_NaT_ conductance values were calculated from peak I_NaT_ measurements using the sodium reversal potential (see Methods). A plot of the mean normalized peak Nav conductance (G/Gmax) against the respective depolarizing voltage step also indicates the voltage-dependence of Nav channel activation is similar in control and *Tsc1*^mut/mut^ Purkinje neurons (Fig. 3D). The kinetics of I_NaT_ fast-inactivation was measured by fitting I_NaT_ decay with a first-order exponential function across voltage-steps. In Figure 3E, the mean (± SEM) time constants (τ) from these fits are plotted for each of the depolarizing voltage-steps, revealing no significant difference between control and *Tsc1^mut/mut^* cells. To measure the voltage-dependence of I_NaT_ steady-state inactivation, varying (100 msec) conditioning voltage steps were applied prior to a common (−20 mV) voltage step used to evoke and measure the proportion of I_NaT_ available for activation (Fig. 3F). In Figure 3G, the mean (± SEM) peak I_NaT_ value, normalized to maximal I_NaT_ (across voltages) for each cell, is plotted against the associated conditioning voltage step. From these plots, no difference was measured in the voltage-dependence of I_NaT_ steady-state inactivation between control and *Tsc1^mut/mut^* cells. Importantly, these Nav current properties were measured in neonatal (P15-16) Purkinje neurons. Using crosses with the Ai14 Cre-reporter line (see Methods) we established Cre-recombinase driven by the L7/Pcp2 promotor is robustly expressed in P15-16 Purkinje neurons (Fig. 1 *A1*). Conversely, in P9 Pcp2-Cre-positive animals, Cre-mediated tdTomato expression is not yet evident (data not shown), indicating the effects of Cre-mediated *Tsc1* deletion, especially as it relates to expression of ion channel proteins, may not yet occur in neonatal Purkinje neurons.

**Figure 3.**
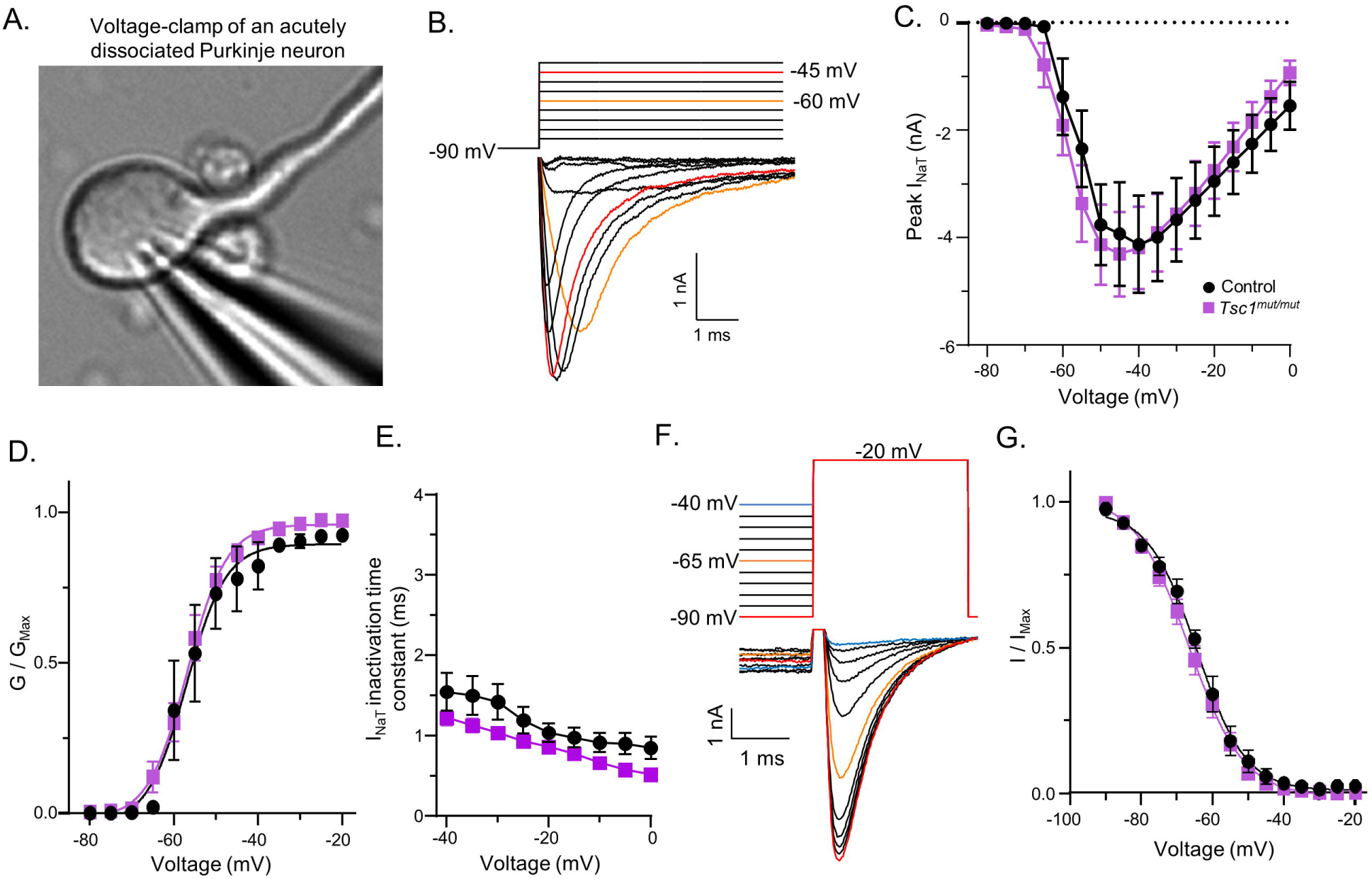
Nav currents are similar in neonatal *Tsc1^mut/mut^* and control Purkinje neurons. In panel **A.,** a representative micrograph of a dissociated neonatal (P15) Purkinje neuron with a glass microelectrode used for patch recording is shown. **B.** Representative voltage-clamp records of evoked I_Na_ during various depolarizing voltage steps are shown from a neonatal control Purkinje neuron. Voltage commands are presented above the current records in corresponding colors. **C.** The mean (± SEM) peak transient sodium current (I_NaT_) is plotted against voltage for control (N = 5, n = 9; *black circles*) and *Tsc1^mut/mut^* (N = 3, n = 11; *magenta squares*) cells and reveals no difference in the amplitudes of peak I_NaT_ evoked across depolarizing voltage steps (RM two-way ANOVA). Similarly, in panel **D.**, plots of the mean (± SEM) normalized conductance values (G/G_Max_) against voltage indicate control and *Tsc1^mut/mut^* neonatal cells have similar voltage dependence of activation (RM two-way ANOVA). Boltzmann fits of control and *Tsc1^mut/mut^* normalized (mean) conductance values have V_1/2_ values of −57 mV and −56.9 mV, respectively. The mean (± SEM) time constant of I_NaT_ inactivation (τ; see Methods) is plotted against voltage in **E.** RM two-way ANOVA analysis of these values revealed no significant difference across genotypes. **F.** The voltage-dependence of I_NaT_ steady-state inactivation was assessed by initially stepping cells to various conditioning voltages before stepping cells to a common (−20 mV) test potential in which peak evoked I_NaT_ was measured. In the representative trace, command voltages are shown above the current trace records in corresponding colors. **G.** Mean (± SEM) normalized I_NaT_ (I/I_Max_) values, measured during the common −20 mV voltage step, are plotted against the preceding conditioning voltage for control (N = 4, n = 9, *black circles*) and *Tsc1^mut/mut^*(N = 3, n = 11, *magenta squares*) cells, revealing no difference in the voltage-dependence of I_NaT_ steady-state inactivation (RM two-way ANOVA). Boltzmann fits of control and *Tsc1^mut/mut^* mean I/I_Max_ plots have V_1/2_ values of −64.4 mV and −66.8 mV, respectively.

Because measurements of attenuated action potential firing in *Tsc1^mut/mut^* Purkinje neurons were acquired from animals 5-8 weeks old, we were interested in testing if *Tsc1* deletion drives changes in Nav current properties from adult Purkinje neurons. The methods used for Purkinje neuron dissociation are not successful in adult (>P21) animals, and so to measure Nav currents from adult (5-8 week-old) Purkinje neurons, we performed voltage-clamp experiments on intact Purkinje neurons in acutely isolated cerebellar slices (Fig. 4A). In these voltage-clamp studies, to limit the contamination of Nav current records by escaped sodium-mediated spikes from distal neurites, we used a pre-pulse protocol that was first developed and utilized by Milescu, Bean, and Smith (2010), and that has been successfully used for Nav measurements in adult Purkinje neurons (Bosch et al., 2015). With these voltage-clamp protocols, membranes were first depolarized to drive Nav channels into the fast-inactivated kinetic state. A subsequent brief repolarization step was used to enable the recovery of a portion of the inactivated Nav channels near the electrode, finally a third voltage-step was used to evoke and measure Nav currents under appropriate space-clamp. These pre-pulse voltage-command protocols (also described in Methods) were used to measure the voltage-dependence of I_NaT_ activation (Fig. 4B) and I_NaT_ steady-state inactivation. Between *Tsc1^mut/mut^* and control Purkinje neurons, we measured no differences in the voltage-dependence of I_NaT_ activation, which is evident from Figure 4C, which shows the mean normalized Nav conductances measured at various test voltages in control and *Tsc1^mut/mut^* Purkinje neurons. Interestingly, however, the mean (±SEM) peak I_NaT_ measured during these depolarizing voltage steps was found to be significantly lower in *Tsc1^mut/mut^* cells compared to control cells (Fig. 4D). Similar to the voltage-clamp results from dissociated Purkinje neurons, no differences were measured in the voltage-dependence of I_NaT_ steady-state inactivation (Fig. 4E) or in the kinetics of I_NaT_ decay (Fig. 4F). Together, these data indicate I_NaT_ gating properties are similar in control and *Tsc1^mut/mut^* adult Purkinje neurons, however, I_NaT_ peak amplitudes are significantly reduced in *Tsc1^mut/mut^* cells, suggesting *Tsc1* deletion may affect the expression and/or localization of Nav channels in *Tsc1^mut/mut^* cells.

**Figure 4.**
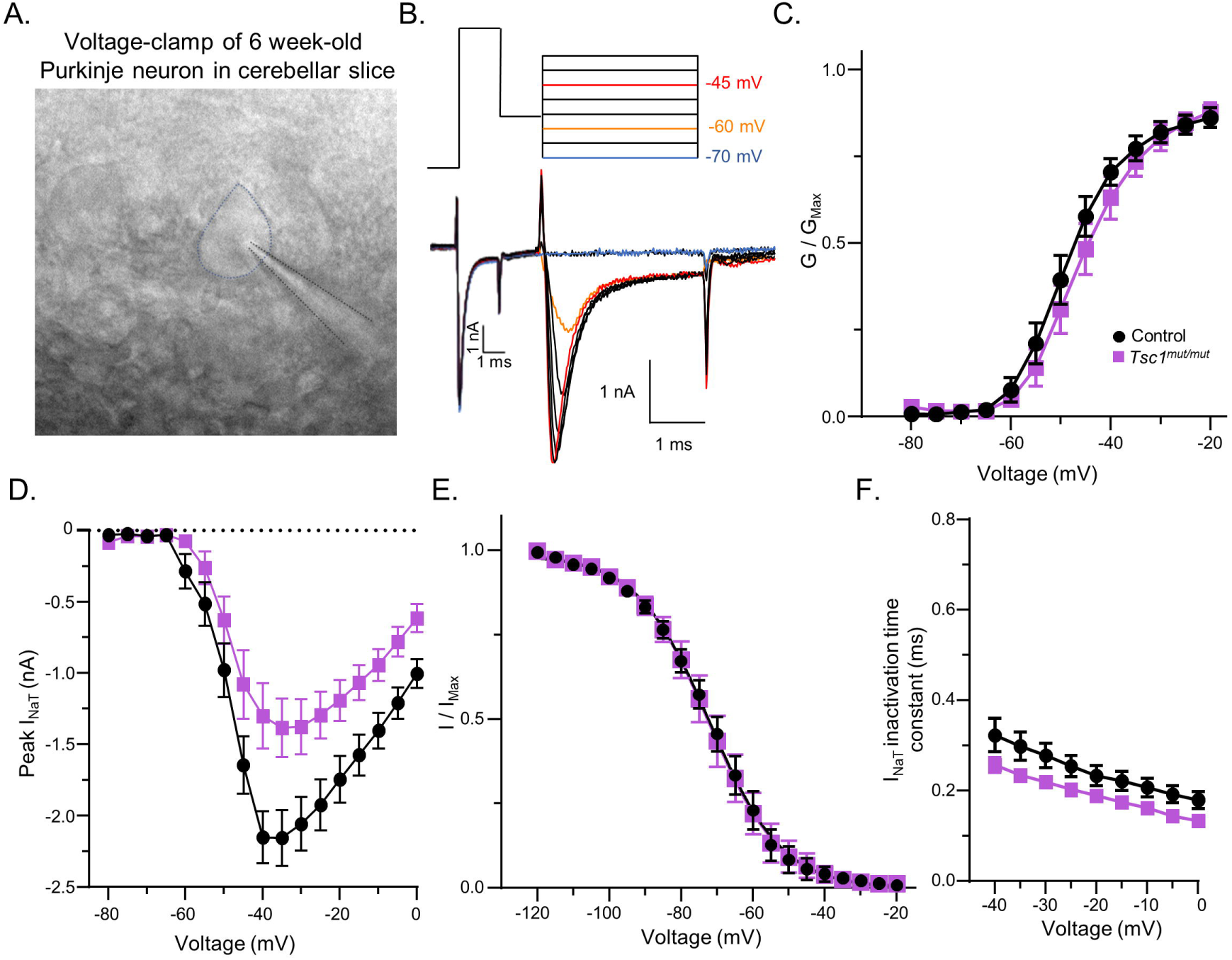
Peak Nav currents are reduced in adult *Tsc1^mut/mut^* Purkinje neurons. **A.** Adult Purkinje neurons were recorded in acutely isolated parasagittal cerebellar slices. In the micrograph, a microelectrode (outlined in black) is shown patching a Purkinje neuron (outlined in blue). **B.** To measure Nav currents in intact Purkinje neurons, a pre-pulse voltage protocol (see Methods) was used measure Nav current properties in the soma and proximal neurite. Representative Nav currents evoked using a protocol to measure the voltage dependence of I_NaT_ activation are shown with the command voltage steps shown above the evoked current traces in corresponding colors. **C.** The normalized peak Nav conductance (G/G_Max_) plotted against activation voltage reveals the voltage-dependence of Nav conductance activation is similar in control and *Tsc1^mut/mut^* Purkinje neurons. Boltzmann fits of control and *Tsc1^mut/mut^* mean conductance values have V_1/2_ values of −48 mV and −45.9 mV, respectively. **D.** The mean (± SEM) peak I_NaT_ values plotted against voltage, however, are significantly (P = .007, RM two-way ANOVA) reduced in *Tsc1^mut/mut^* cells compared to control cells (control N = 6, n = 22, *black circles*; *Tsc1^mut/mut^* N = 6, n = 16, *magenta squares*). **E.** Plots of the normalized peak I_NaT_ (I/I_Max_) evoked at a common −20 mV voltage step, against the conditioning voltage (see Figure 3F) reveal no change in the voltage-dependence of I_NaT_ steady-state inactivation (control: N = 5, n = 18; *black circles*; *Tsc1^mut/mut^*: N = 5, n = 13; *magenta squares*). Boltzmann fits of control and *Tsc1^mut/mut^* mean I/I_Max_ plots have V_1/2_ values of −72.1 mV and −72.6 mV, respectively. **F.** A plot of the time constant of I_NaT_ inactivation (τ; see Methods) against the activating voltage step reveals control (N = 6, n = 22; *black circles*) and *Tsc1^mut/mut^*(N = 6, n = 16; *magenta squares*) Purkinje neuron I_NaT_ inactivation kinetics are not significantly different (RM two-way ANOVA).

### Integrated anti-ankyrinG and anti-pan Nav immunofluorescence are diminished at the AIS of *Tsc1^mut/mut^* Purkinje neurons

To explore if the reduced amplitude of peak I_NaT_ in *Tsc1^mut/mut^* cells (Fig. 4D) reflects a change in the expression properties of Nav channels, secondary immunofluorescence was used to label Nav channels expressed along the Purkinje neuron AIS. AnkyrinG secondary immunofluorescence was used as a marker for the Purkinje neuron AIS. Calbindin immunofluorescence or Cre-mediated tdTomato expression (in Ai14 crossed mice, see Methods), were used to identify immunolabeling on membranes specific to Purkinje neurons. In Figure 5A, immunolabeling in a parasagittal cerebellar section from a Cre-positive control animal (crossed with the Ai14 Cre-reporter line) is shown. Anti-pan Nav channel immunofluorescence (shown in green, panel 2) is co-localized with anti-ankyrinG fluorescence (*red*, panel 3) along the AIS segment extending from Purkinje neuron somata (*magenta,* panel 4). The combined image is shown in panel 1 of Figure 5A with co-localized anti-pan Nav and anti-ankyrinG immunofluorescence along Purkinje neuron AISs appearing yellow. In Figure 5B, the fluorescence signal of pan-Nav and anti-ankyrinG secondary immunolabeling are compared between the AIS of an adult Purkinje neuron from a control animal (upper panels) and a *Tsc1^mut/mut^* animal (lower panels). Using line-scans drawn along the AIS region extending from the Purkinje neuron soma, we measured the intensity values of anti-pan Nav and anti-ankyrinG immunofluorescence between control and *Tsc1^mut/mut^* animals. The mean (±SEM) intensity values for anti-ankyrinG (Fig. 5C) and anti-pan Nav (Fig. 5D) immunofluorescence are clearly diminished along the AIS of *Tsc1^mut/mut^* cells compared to controls. From these plots of fluorescence intensity along the AIS of cells from control and *Tsc1^mut/mut^* animals, we calculated the area under the curve values (integrated fluorescence intensity along the AIS), which revealed significantly reduced fluorescence for anti-ankyrinG (Fig. 5E) and anti-pan Nav (Fig. 5F) labeling in *Tsc1^mut/mut^* cells.

**Figure 5.**
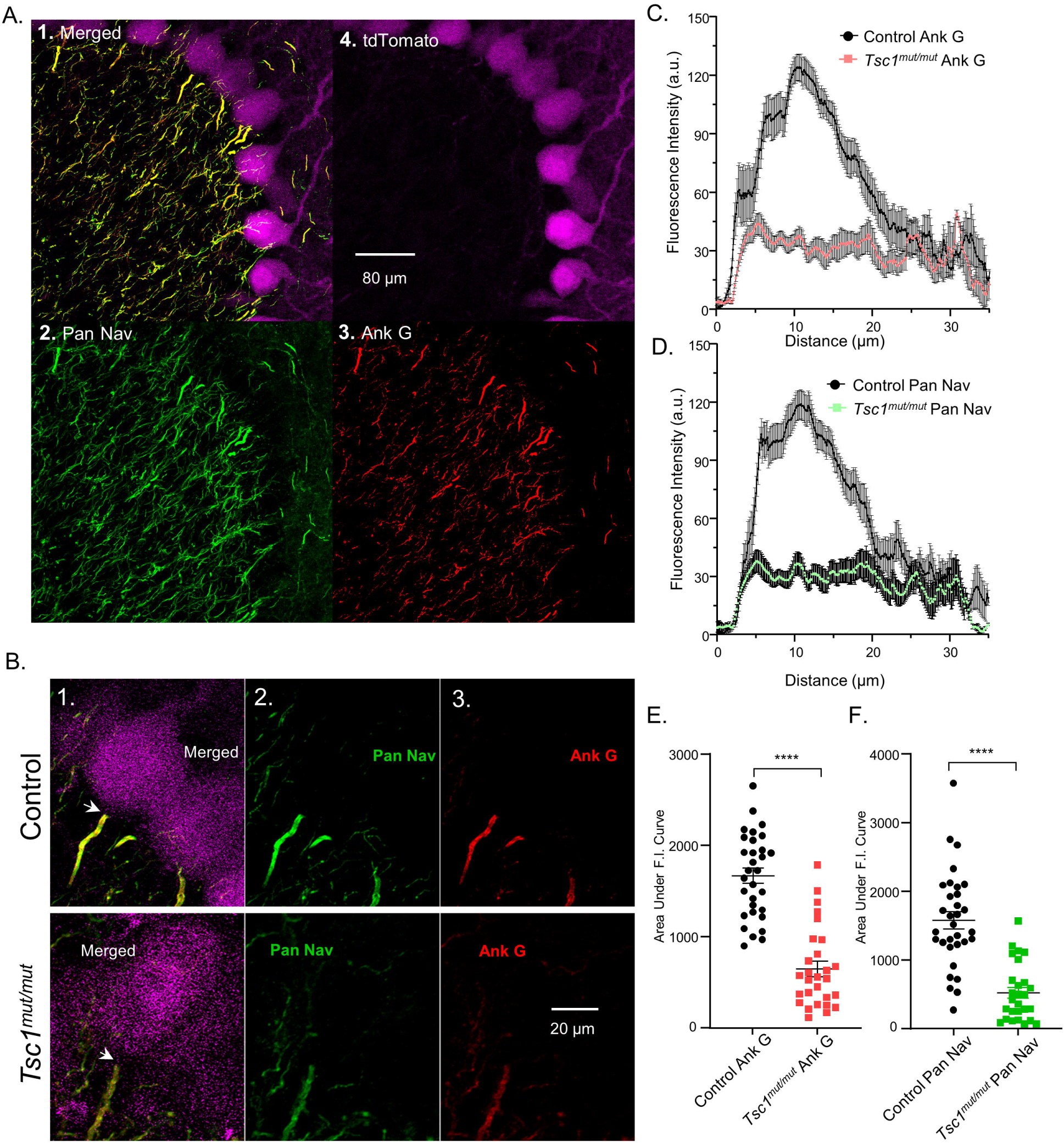
Anti-pan Nav and anti-ankyrinG immunofluorescence is reduced at the axon initial segment of *Tsc1^mut/mut^* Purkinje neurons. **A.** Secondary antibody fluorescent labeling was used to localize and measure the intensity of anti-pan Nav (A. 2.) and anti-ankyrinG (A. 3.) labeling in Purkinje neurons labeled by Cre-dependent tdTomato expression (A. 4.) shown in *magenta*. Combined images are shown in image A.1. with anti-pan Nav (*green*) and anti-ankyrinG (*red*) dual labeling at Purkinje neuron AIS appearing *yellow*. Images in A. were acquired from an adult control animal. In panel **B.**, anti-pan Nav and anti-ankyrinG labeling are presented and compared directly between individual Purkinje neurons from control (*upper*) and *Tsc1^mut/mut^* (*lower*) animals with merged (B. 1.), single channel anti-pan Nav (B. 2.), and single channel anti-ankyrinG (B. 3.) presented in panels 1, 2, and 3. The mean (± SEM) intensity values of anti-ankyrinG **(C.)** and anti-pan Nav **(D.)** immunofluorescence are plotted against distance along the AIS of control (*black circles*) and *Tsc1^mut/mut^* (*squares*) Purkinje neurons. Line scans were used to localize intensity measurements to the AIS (see Methods). For intensity measures along the AIS of each cell, an area under the curve (AUC) value was used to determine the integrated fluorescence intensity of the anti-pan Nav or anti-ankyrinG signal at the cell AIS. Plots of these values reveal significantly (P < .0001, Student’s unpaired t-test) reduced anti-ankyrinG (**E.**) and anti-pan Nav (**F.**) signals at the AIS of *Tsc1^mut/mut^* Purkinje neurons (N = 4, n = 28) compared to Purkinje neurons from control animals (N = 4, n = 31).

### Heterozygous *Tsc1* deletion causes slight changes in AIS immunofluorescence and does not affect membrane excitability

We tested if the selective deletion of a single *Tsc1* allele also affects the integrated anti-ankyrinG and anti-pan Nav immunofluorescence intensity along the AIS. In these experiments, comparing AIS labeling in control and heterozygous (*Tsc1^mut/+^*;Cre-positive) Purkinje neurons, we measured significant reductions in the mean (±SEM) integrated intensity of both anti-ankyrinG (Fig. 6A, B) and anti-pan Nav (Fig. 6C, D) labeling (P < .01, Welch’s unpaired t-test). Mean (±SEM) anti-ankyrinG and anti-pan Nav channel integrated AIS immunofluorescence from *Tsc1^mut/+^* cells was also found to be significantly (P < .0001, Welch’s unpaired t-test) higher than measurements from *Tsc1^mut/mut^* cells (plotted in Fig. 5E, F).

**Figure 6.**
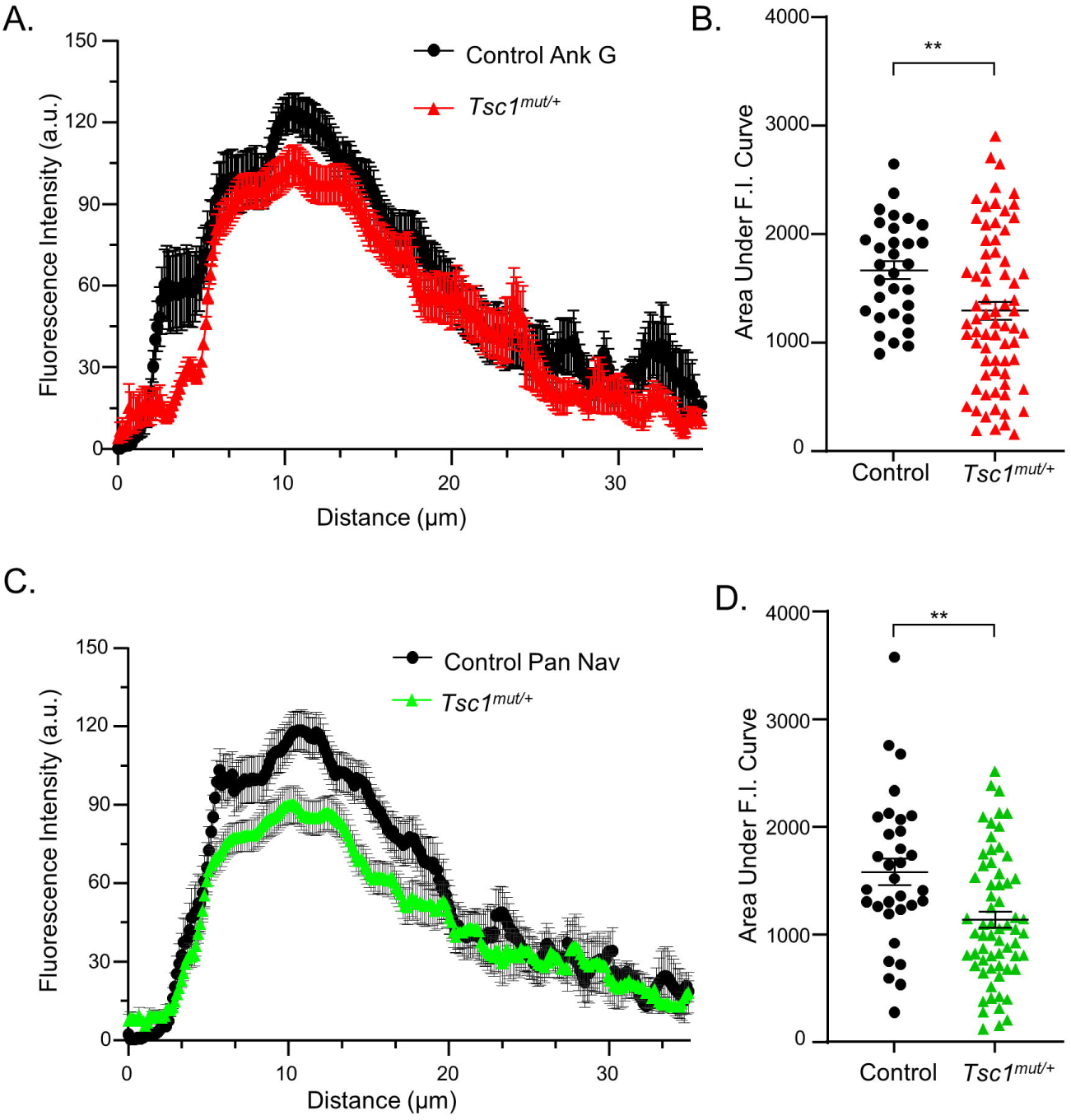
Heterozygous *Tsc1* deletion (*Tsc1^mut/+^*) results in slight reductions in anti-ankyrinG and anti-pan Nav labeling at the Purkinje neuron AIS. **A.** The mean (± SEM) fluorescence intensity values of anti-ankyrinG (**A.**) and anti-pan Nav (**B.**) immunolabeling along the AIS of *Tsc1^mut/+^* Purkinje neurons (N = 3, n = 64) were compared to intensity values from control Purkinje neurons (N = 4, n = 31). Analyses of these data via area under the curve (AUC) measurements for individual cells revealed slight, but significantly lower (Welch’s unpaired t-test, P < .01) integrated fluorescence intensity of anti-ankyrinG (**C.**) and anti-pan Nav (**D.**) labeling along the AIS of *Tsc1^mut/+^* Purkinje neurons compared to control cells.

We tested if *Tsc1^mut/+^* Purkinje neurons have deficits in repetitive firing (representative firing shown in Fig. 7A) that, similar to the immunofluorescence measurements, are also intermediate to the firing properties measured in wild type control and *Tsc1^mut/mut^* (Figure 2) Purkinje neurons. Interestingly, *Tsc1^mut/+^* Purkinje neurons share similar measures of intrinsic excitability as wild type control Purkinje neurons. In Figure 7, the mean (±SEM) spontaneous and evoked repetitive firing frequencies are plotted for controls and *Tsc1^mut/+^* Purkinje neurons (Fig 7B, C), revealing no significant differences. Additionally, action potential waveform measurements, including the action potential threshold voltage, APD, and action potential amplitudes (Fig. 7D, E, F), are also similar between these groups.

**Figure 7.**
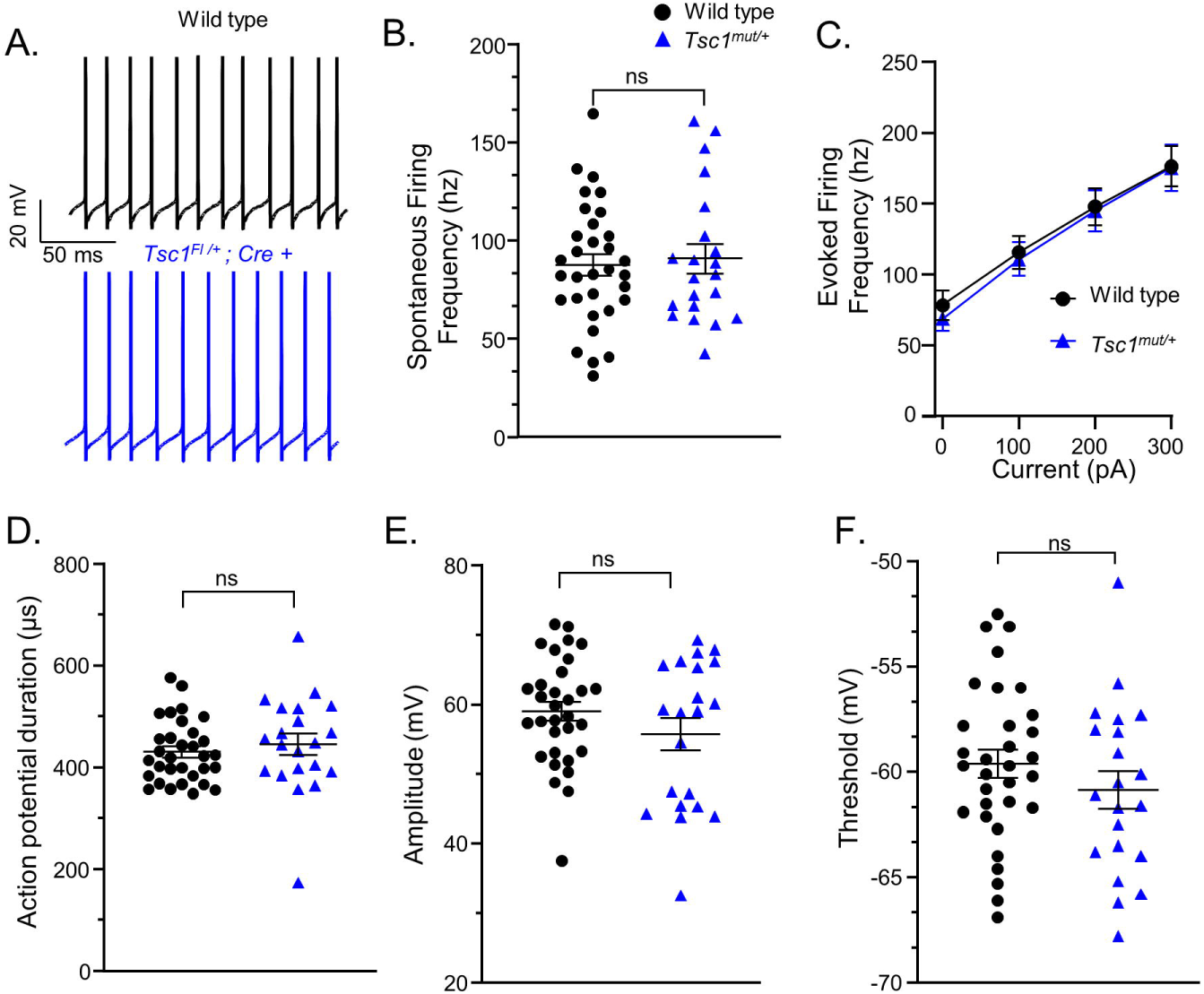
Heterozygous *Tsc1* deletion (*Tsc1^mut/+^*) does not affect Purkinje neuron firing. **A.** Spontaneous repetitive firing was recorded in 5-8 week-old wild type control (*black*) and *Tsc1^mut/+^*(blue) Purkinje neurons. Measurements of repetitive firing during spontaneous activity, plotted in **B.** (unpaired Student’s t-test), or during depolarizing current injections, plotted in **C.** (RM two-way ANOVA), reveal the deletion of a single *Tsc1* allele does not significantly affect spontaneous and evoked firing frequency. Similarly, action potential waveform properties of *Tsc1^mut/+^* Purkinje neurons, including the action potential duration (**D.**), amplitude (**E.**), and action potential threshold voltage (**F.**), are not significantly different (unpaired Student’s t-test) from measurements in wild type control cells (wild type: N = 13, n = 32; *Tsc1^mut/+^*: N = 6, n = 21).

### Action potential derivative plots reveal action potential initiation at the AIS is impaired in *Tsc1^mut/mut^* Purkinje neurons

The AIS in Purkinje neurons functions as the site of action potential generation/initiation (Khaliq and Raman, 2006). Purkinje neuron action potentials propagate to vestibular and deep cerebellar nuclei, driving GABA release onto post-synaptic cells. Action potentials also backpropagate into the Purkinje neuron soma causing a secondary (somatic) spike waveform. AIS (primary) and somatic (secondary) spikes can be resolved through analysis of the first and second derivative of the action potential waveforms (Stuart and Häusser, 1994; Bean, 2007; Meeks and Mennerick, 2007). Because our findings suggest *Tsc1* deletion affects Purkinje neuron firing properties via Nav channel localization at the AIS, we examined 2^nd^ derivative plots of action potential waveforms from *Tsc1^mut/mut^* and control Purkinje neurons to examine if spike initiation and backpropagation into the somatic compartment are affected. Representative action potentials (Fig. 8A), with the corresponding 1^st^ (*A1*, dV/dt) and 2^nd^ (*A2*, dV^2^/dt) derivative voltage plots (time-locked with action potential traces), are shown for control and *Tsc1^mut/mut^* current-clamp records. Notably, in *Tsc1^mut/mut^* cells, the 1^st^ derivative plot over time (dV/dt) reveals a pronounced hitch, (highlighted by the green arrow) during the upstroke of the curve. In the *Tsc1^mut/mut^* 2^nd^ derivative record, this hitch in the 1^st^ derivative plot corresponds with a clear separation between positive peaks that reflect spike initiation in the AIS (first peak) and in the somatic compartment (second peak). Asterisks highlight the positive peaks in the second derivative voltage plots (*A2*). Compared to similar plots from a control Purkinje neuron, this difference reveals an increase in the delay between action potential generation at the AIS and in the soma. The delay between spike generation is quantified by measuring the time between the two peaks of the the 2^nd^ derivative plots. The mean ± SEM delay between AIS and somatic spikes is significantly longer in *Tsc1^mut/mut^* Purkinje neurons compared to wild type controls (Fig. 8B), suggesting deficits in the back propagation of spikes generated in the AIS into the somatic compartment of these neurons. It is also evident from the first derivative representative plots (Fig. 8 *A1*) that the maximal dV/dt is lower in *Tsc1^mut/mut^* cells. This effect is directly evident in phase plots in which dV/dt is plotted against voltage for *Tsc1^mut/mut^* and control cells (shown in Fig. 8C). In Figure 8D, the peak dV/dt values for action potentials recorded from wild type control and *Tsc1^mut/mut^* cells are plotted, revealing *Tsc1^mut/mut^* Purkinje neurons indeed have significantly lower mean (± SEM) peak dV/dt values, which is consistent with the reduced I_NaT_ and anti-pan Nav labeling measured at the AIS of *Tsc1^mut/mut^* Purkinje neurons. Longer APD in *Tsc1^mut/mut^* cells (presented in Fig. 2D) also corresponds with action potentials from these cells having significantly reduced slope values (compared to control cells) during the action potential upstroke (measured as maximum dV/dt, Fig. 8D, Table 1). Notably, minimum dV/dt, reflective of action potential downstroke slope, is also significantly reduced in *Tsc1^mut/mut^* cells (see Table 1), which may reflect changes in the expression and/or gating properties of other ionic currents/channels necessary for action potential termination and/or changes in membrane passive properties.

**Figure 8.**
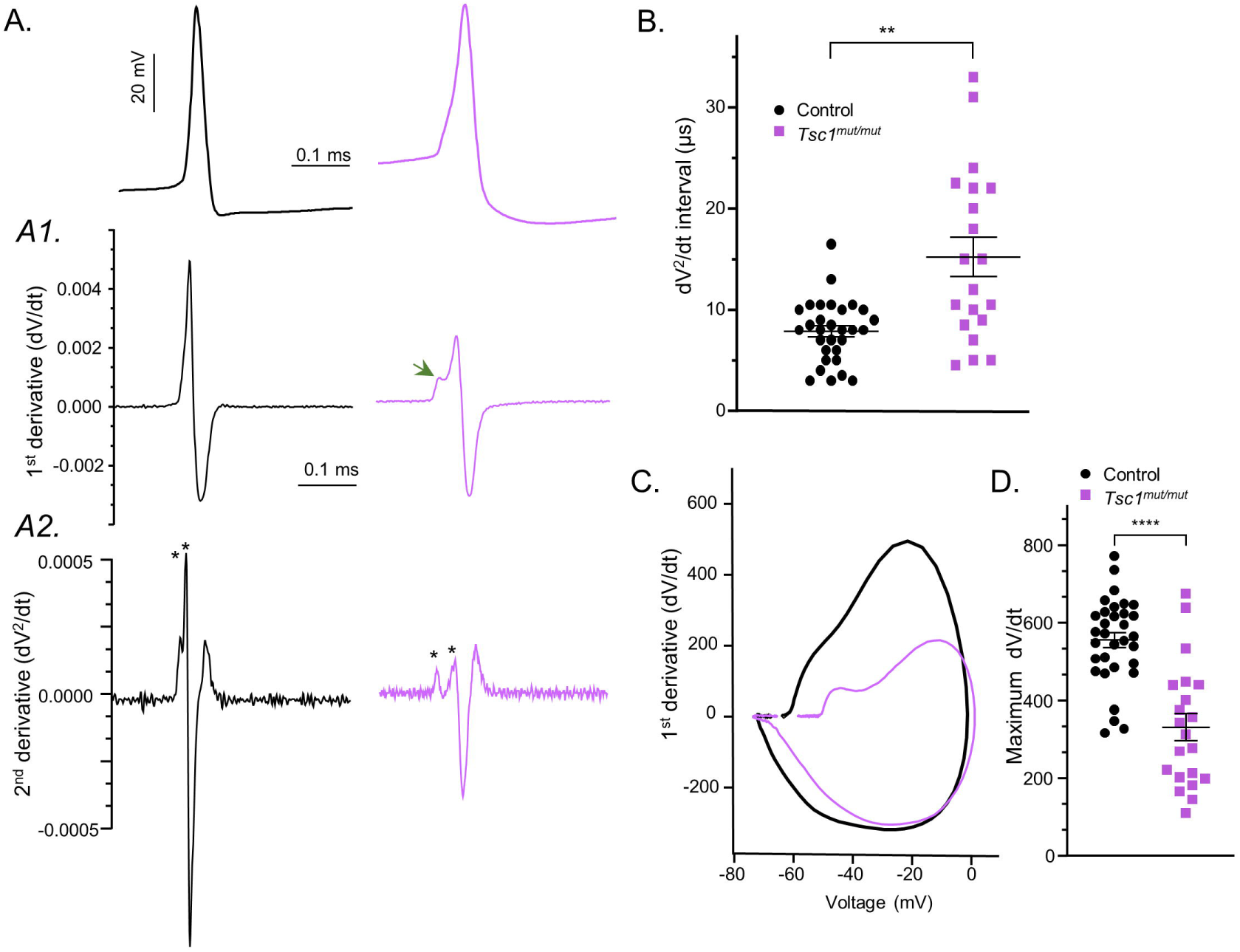
Action potential derivative analyses reveal altered action potential initiation and propagation in *Tsc1^mut/mut^* Purkinje neurons. **A.** Representative action potential waveforms are presented from 6 week-old control (*black*) and *Tsc1^mut/mut^* (*magenta*) Purkinje neurons. In the *A1* and *A2* panels (*below*), plots time-locked with the control and *Tsc1^mut/mut^* action potentials show the corresponding voltage 1^st^-derivative (dV/dt, *A1*) and voltage 2^nd^-derivative (dV^2^/dt, *A2*). The green arrow highlights the hitch in the dV/dt plot during the action potential upstroke. Asterisks in *A2* denote spike initiation in the AIS (first) and after a short delay, in the somatic compartment (see Methods). **B.** Across cells, this delay, measured using 2^nd^-derivative plots, was determined to be significantly longer in *Tsc1^mut/mut^* cells compared to wild type controls (Welch’s unpaired t-test, P = .0012; wild type controls: N = 13, n = 32; *Tsc1^mut/mut^*: N = 6, n = 21). In **C.**, representative phase plots corresponding to the action potentials presented in panel A. are shown. **D.** The maximum dV/dt during the action potential upstroke is significantly (unpaired Student’s t-test, P < .0001) reduced *Tsc1^mut/mut^* cells compared to wild type controls.

### Partial Nav channel block recapitulates effects of *Tsc1* deletion on Purkinje neuron action potential waveforms

Nav channels expressed along the AIS are the site of action potential generation and regulate the repetitive firing frequencies in cerebellar Purkinje neurons (Khaliq and Raman, 2006; Bosch et al., 2015). Homozygous *Tsc1* deletion in Purkinje neurons results in reduced anti-pan Nav labeling at the AIS, which corresponds to attenuated repetitive firing frequency and changes in the action potential waveform. To examine if Nav channels are directly responsible for these changes, we used subsaturating concentrations of the selective Nav channel blocker TTX to test how partial Nav channel block/loss affects action potential properties in control Purkinje neurons. Partial Nav channel block resulted in similar changes to the action potential waveform as those measured in *Tsc1^mut/mut^* cells (Fig. 9A). Across six wild type cells, 1 nM TTX resulted in a significant depolarized shift in the action potential threshold voltage (Fig. 9B), an increase in APD (Fig. 9C), and reduced action potential amplitude (Fig. 9D). Additionally, the maximum dV/dt in the action potential waveforms were significantly reduced after TTX exposure (Fig. 9E) and the delay between spike initiation in the AIS and somatic compartment, measured from 2^nd^ derivative action potential plots, was significantly increased (Fig. 9F) after TTX exposure. These changes in the action potential waveform after partial Nav channel block suggest reduced Nav channel expression/availability at the AIS of *Tsc1^mut/mut^* cells is a major contributor to the impaired excitability of these cells. It’s important to note, however, that partial Nav channel block (under 1 nM TTX) causes Purkinje neurons to transition from a state of sustained repetitive firing to exhibiting intermittent quiescence, which is followed by several seconds repetitive action potential firing that often includes intermittent calcium spikes (data not shown). This effect of TTX on Purkinje neuron firing is a clear deviation from the reduced repetitive action potential firing measured in *Tsc1^mut/mut^* Purkinje neurons, which are sustained and lack small amplitude calcium spikes. These differences likely reflect that *Tsc1* deletion drives additional changes, beyond reduced Nav channel expression, that impact Purkinje neuron excitability. These differences may also reflect that TTX exposure causes a sudden loss of Nav channels/conductance without an opportunity for compensatory mechanisms to restore/maintain the sustained repetitive firing measured in *Tsc1^mut/mut^* cells.

**Figure 9.**
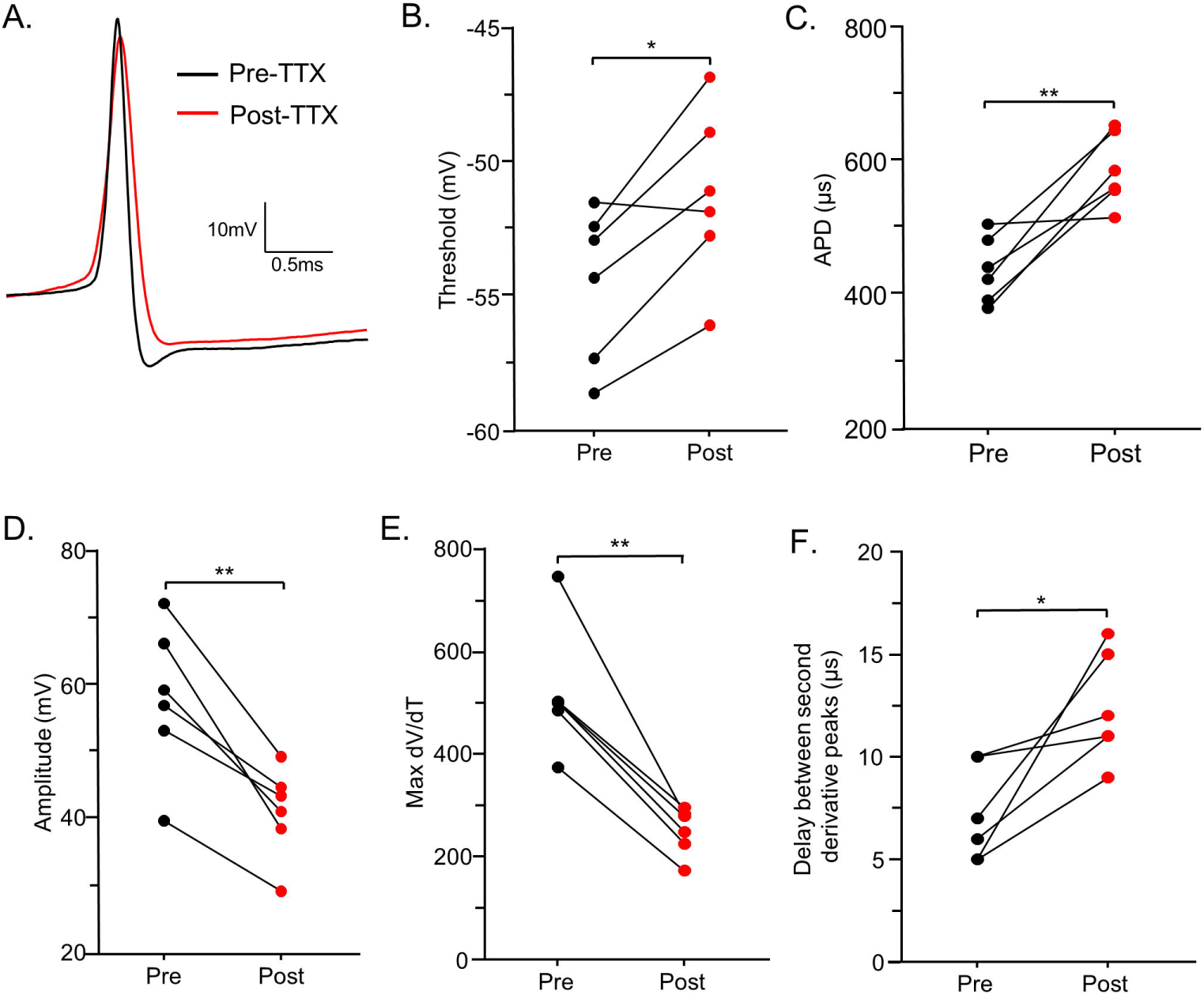
Partial Nav channel block causes similar changes to the action potential waveform as those measured in *Tsc1^mut/mut^* cells. **A.** Measures of spontaneous action potential were performed in control Purkinje neurons before (*black*) and after (*red*) applying 1 nM tetrodotoxin (TTX). TTX is a selective Nav channel blocker and 1 nM TTX partially blocks Purkinje neuron Nav channels resulting in significant changes to the action potential waveform. **B.** Changes include a significant (P < .05) depolarizing increase in the action potential threshold voltage, (**C.**) a significant increase in the action potential duration (P < .01), (**D.**) and a significantly reduced action potential amplitude (P < .01). **E. – F.** Analyses of action potential 1^st^ and 2^nd^ derivatives revealed TTX exposure significantly (P < .01) reduces peak dV/dt and increases (P < .01) the delay between the second derivative peaks. Paired Student’s t-tests were used for pre- and post-TTX comparisons. Experiments were performed on 6 Purkinje neurons in cerebellar slices from two (N = 2) 6 week-old wild type animals.

## DISCUSSION

Deletion of *Tsc1* selectively in cerebellar Purkinje neurons results in multiple ASD-like behavioral phenotypes that are linked to attenuated Purkinje neuron firing (Tsai et al., 2012, 2018; Lawson et al., 2024). TSC1 and TSC2, known as hamartin and tuberin, respectively, combine to form a protein complex that acts as a GTPase activating protein (GAP) for the small GTPase Rheb, effectively inhibiting its activity and subsequently downregulating the mTORC1 pathway (Kwiatkowski, 2003). The diverse and varying symptoms linked to tuberous sclerosis are driven by global autosomal dominant loss-of-function mutations in *TSC1* or *TSC2* (European Chromosome 16 Tuberous Sclerosis Consortium, 1993; Crino et al., 2006; Au et al., 2007), which are likely to cause different and potentially more numerous deficits in Purkinje neuron and cerebellar circuit function than those measured in the *Tsc1* mutant model utilized here. Across cells, however, inactivation of either *Tsc1* or *Tsc2* causes exaggerated mTORC1 signaling and excessive cellular growth and metabolism (Wullschleger et al., 2006; Inoki and Guan, 2009), which can, over time, drive stress-mediated apoptosis (Ng et al., 2011). Purkinje neurons in the *Tsc1^mut/mut^* model are no exception. *Tsc1^mut/mut^* Purkinje neurons have increases in soma size, greater numbers of dendritic spines, and reduced survival in animals ≥8 weeks-old (Tsai et al., 2012). In younger (6 week-old) *Tsc1^mut/mut^* animals, as well as in heterozygote *Tsc1* deletion animals (*Tsc1^mut/+^*), no significant changes in Purkinje neuron survival were detected (Tsai et al., 2012). Membrane capacitance is a common measure of membrane surface area (Golowasch et al., 2009). In our current-clamp recordings from mutant (*Tsc1^mut/mut^* and *Tsc1^mut/+^*) Purkinje neurons, average membrane capacitance values were higher than wild type control cells, but not significantly different (Table 1). In *Tsc1^mut/mut^* cells, taken from 5-7 week-old animals, we measured significantly attenuated repetitive firing frequency compared to wild type controls, suggesting the ASD-related behavioral phenotypes previously measured in these *Tsc1^mut/mut^* animals are at least initially, a result of attenuated Purkinje neuron firing. In these current-clamp studies, we did not find mean firing frequencies to differ between male and female animals or between cells taken from younger (≤6 week-old) and older (>6 week-old) animals ((within both wild type and *Tsc1^mut/mut^* datasets, see Supplemental Figure 1).

We investigated the biophysical/molecular mechanisms underlying *Tsc1^mut/mut^* Purkinje neuron deficits in repetitive firing and uncovered diminished peak Nav current and reduced secondary fluorescent labeling of Nav channels at the axon initial segment (AIS) of adult *Tsc1^mut/mut^* cells. Using subsaturating concentrations of the selective Nav channel blocker TTX, we determined an acute partial block of Nav channels in wild type Purkinje neurons results in changes in the action potential waveform that mirror those measured in *Tsc1^mut/mut^* cells. The AIS has previously been shown to be the site of action potential initiation and is critical for regulating repetitive simple spike activity in mouse Purkinje neurons (Khaliq and Raman, 2006; Bosch et al., 2015). In *Tsc1^mut/mut^* Purkinje neurons, we found an increased delay in the propagation of action potentials from the AIS into the somatic compartment. Interestingly, ankyrinG, a cytoskeletal linker protein and critical organizer of the AIS, was also found to have reduced secondary immunofluorescence at the AIS of *Tsc1^mut/mut^* Purkinje neurons, suggesting the reduced Nav channels/currents may reflect a more general dysregulation and function of the AIS in *Tsc1^mut/mut^* cells.

### Molecular mechanisms of impaired firing in *Tsc1* mutants

Tuberous sclerosis is an autosomal dominant disorder caused by global loss-of-function mutations in *TSC1* or *TSC2* (Northrup, 1992). In the presented data, we detected no changes in the intrinsic excitability of *Tsc1^mut/+^* Purkinje neurons from 5-8 week-old mice, and significant (although slight) reductions in anti-pan Nav and anti-ankyrinG labeling along the AIS of *Tsc1^mut/+^* animals. Tsai et al. (2012) previously measured attenuated spontaneous and evoked firing frequencies of *Tsc1^mut/+^* cells from 4-6 week-old animals, and these changes in excitability corresponded with deficits in social interaction behavior (measured in 7-9 week-old mice), but interestingly, no deficits in motor coordination (assessed via the rotarod assay) or in gait (Tsai et al., 2012). In a previous investigation, we also determined balance and motor coordination were not impaired in 9-11 week-old *Tsc1^mut/+^* animals, while social interaction deficits were measured in 9-11 week-old *Tsc1^mut/+^* males (Lawson et al., 2024). Given that we measured significantly reduced anti-pan Nav and anti-ankyrinG AIS immunofluorescence in *Tsc1^mut/+^* cells, it is surprising we did not measure a firing deficit in 5-8 week-old *Tsc1^mut/+^* Purkinje neurons. It’s possible changes driven by *Tsc1* haploinsufficiency in Purkinje neurons are progressive and will eventually cause AIS dysfunction in older *Tsc1^mut/+^* cells. Our measurements of firing and AIS organization were confined to Purkinje neurons within lobules II-VI of the spinocerebellum (vermis and paravermis) and so it is also possible that Purkinje neurons in the cerebellar hemispheres are differentially affected by heterozygous *Tsc1* deletion.

In these studies, the attenuated firing of *Tsc1^mut/mut^* Purkinje neurons was linked to impaired initiation and propagation of action potentials at the AIS, and reduced anti-ankyrinG and anti-pan Nav integrated immunofluorescence at the Purkinje neuron AIS. However, other factors may also contribute to this reduced excitability. We found *Tsc1^mut/mut^* cells have significantly lower input resistance compared to control cells (Table 1, Tsai et al., 2012), which may contribute to or cause the reduced maximum and minimum dV/dt values (action potential slope) measured in these cells. While the reported changes in membrane input resistance were taken from the Purkinje neuron somatic compartment, changes in AIS morphology and/or leak channel expression may reduce input resistance within the AIS and directly impact action potential generation. mTORC1 inhibition has previously been shown to cause increases in calcium-activated potassium current (I_KCa_) in CA1 pyramidal neurons (Springer et al., 2014). If this mechanism is conserved in Purkinje neurons, targeted deletion of *Tsc1*, and the resulting increase in mTORC1 activity, may result in reduced I_KCa_ and contribute to the impaired firing of *Tsc1^mut/mut^* cells. Additionally, there may be some level of Purkinje neuron death/apoptosis, even in 6 week-old *Tsc1^mut/mut^* animals. Purkinje neuron axon collaterals have been shown to form synapses on neighboring Purkinje cells in the parasagittal plane (Witter et al., 2016), and thus, a loss of these synapses due to neighboring cell death may affect the firing properties of *Tsc1^mut/mut^* Purkinje neurons. However, because these synapses between Purkinje cells are inhibitory (Ito et al., 1964; Obata et al., 1967), we would not expect a loss of these synaptic inputs to drive reduced membrane excitability.

Similar to its role in AIS structure/function, ankyrinG is thought to be critical for the functional clustering of ion channels and scaffolding proteins at axonal nodes (Dzhashiashvili et al., 2007; Yang et al., 2007; Gasser et al., 2012), however, in peripheral sensory neurons, it was found that selective ankyrinG deletion results in compensatory expression of ankyrin-R and βI-spectrin at nodal junctions, rescuing the clustering of Nav channels at these nodes (Ho et al., 2014). The effect of *Tsc1* deletion on the organization and functioning of axonal nodes has not been investigated in Purkinje neurons (or other neuronal cell types). Voltage-gated sodium and potassium channels are clustered at nodal and paranodal junctions along the axons of Purkinje cells, which is necessary for spike propagation and GABA release onto vestibular and DCN post-synaptic terminals (Barron et al., 2018). Importantly, if there is a loss of Nav channel expression along Purkinje neuron axonal nodes, it would suggest circuit deficits in the *Tsc1^mut/mut^* mouse model might also (or primarily) be driven by the failure of action potential propagation in Purkinje neuron axons.

### Dysregulated axon initial segment in *Tsc1^mut/mut^* Purkinje neurons

AnkyrinG functions as a molecular scaffold that recruits cytoskeletal and channel proteins to the AIS and other neuronal compartments (Yoon et al., 2022). Silencing ankyrinG expression results in the loss of the AIS and causes axons to acquire dendritic characteristics (Hedstrom et al., 2008). The longest isoform of ankyrinG (480 kDa) interacts with end-binding proteins to drive AIS formation and establish neuronal polarity (Fréal et al., 2016). As a functional organizer of the AIS, ankyrinG is essential for recruiting other AIS proteins, including βIV-spectrin and Nav channels. While βIV-spectrin localization depends on its interaction with ankyrinG, disrupting βIV-spectrin does not affect ankyrinG or the ankyrinG-mediated clustering of Nav channels (Yang et al., 2007). We also know targeted deletion of *Fgf14*, an accessory subunit which binds and regulates Nav channels, as well as targeted deletion of *Scn8a*, which encodes the Nav1.6 pore-forming α subunit, does not affect the expression/localization of ankyrinG at the Purkinje neuron AIS (Xiao et al., 2013; Bosch et al., 2015). These previous reports, which together highlight ankryinG as the primary organizer of the AIS, suggest the diminished anti-pan Nav immunofluorescence at the AIS of *Tsc1^mut/mut^* Purkinje neurons may be directly caused by reduced ankyrinG expression.

Intracellular Fgf14 (iFgf14) interacts with Nav channels, regulating the voltage-dependence of Nav channel steady-state inactivation (Bosch et al., 2015; Ransdell et al., 2024). In the voltage-clamp experiments presented in Figures 3 and 4, we found no changes in Nav channel gating properties, including the voltage-dependence of steady-state inactivation, suggesting iFgf14-mediated regulation, and other mechanisms of post-translational regulation of Nav channel gating remain intact in *Tsc1^mut/mut^* Purkinje neurons. The lack of a change in Nav channel gating was surprising due to the measured depolarized shift in the action potential threshold voltage in *Tsc1^mut/mut^* cells (Fig. 2B), which suggested a potential change in the voltage-dependence of Nav channel activation. Interestingly, application of subsaturating TTX, resulting in the partial and selective block of Nav channels, also resulted in a depolarized shift in action potential threshold voltage, indicating a loss of Nav channels/Nav current can also underlie a depolarized shift in action potential threshold voltage.

Moving forward, to more accurately investigate changes in AIS properties in *Tsc1* mutant Purkinje neurons, super-resolution microscopy techniques such as STORM may provide additional detail, especially if performed across sequential age groups, to delineate if reduced ankyrinG expression at the AIS precedes changes in sodium channel alpha subunit expression. Identifying how these changes correspond with morphological changes, firing properties, and Purkinje neuron survival may also shed light on the primary drivers of Purkinje neuron apoptosis in this model.

It’s not clear why *Tsc1* deletion, and potentially the exaggerated activity of mTORC1, results in ankyrinG dysregulation. To date, there has not been a direct association of mTORC1 signaling with ankyrinG expression/localization. However, ankyrinG at the AIS of primary cortical neurons has been shown to be negatively regulated by activation of the NF-κB transcription factor (König et al., 2017). NF-κB is a ubiquitously expressed transcription factor maintained in its inactive form in the cytosol, and on activation, is translocated into the nucleus (Bonizzi and Karin, 2004). NF-κB activation relies on IκB kinase (IKK) activity (Liu et al., 2012), which has also been demonstrated to phosphorylate and increase mTORC1 activity (Dan et al., 2007, 2014). In a tumor cell model of tuberous sclerosis, increases in mTORC1 activity have been shown to also activate IKK/ NF-κB (Gao et al., 2015). The convergence of these pathways suggests exaggerated mTORC1 activity after *Tsc1* deletion (Tsai et al., 2012) may result in a corresponding increase in the activation of NF-κB, driving reduced ankyrinG expression.

Our data bring into question if *Tsc1* deletion in other neuronal cell types has a conserved deleterious effect on the functioning of the AIS. Across cell types, the loss of *Tsc1* appears to have varying effects on neuronal firing, although results from these studies have typically revolved around changes to synaptic strength/function. For instance, loss of *Tsc1* in hippocampal neurons was found to drive hyperexcitability in hippocampal cultures via reduced synaptic inhibition onto excitatory pyramidal neurons (Bateup et al., 2013). In layer 2/3 cortical pyramidal neurons, *Tsc1* deletion results in reduced inhibitory synaptic currents mediated by GABA receptors but has no effect on excitatory currents. In the striatum, however, *Tsc1* deletion was shown to selectively enhance the intrinsic excitability in striatonigral, but not striatopallidal neurons. The increased excitability of striatonigral cells was associated with significantly reduced inwardly rectifying potassium currents and an increase in membrane input resistance (Benthall et al., 2018). Notably, in *Tsc1^mut/mut^* Purkinje neurons, we found mean input resistance was significantly reduced (Table 1), an effect also reported by Tsai et al. (2012). Interestingly, rheobase current in the *Tsc1* knockout striatonigral neurons was significantly reduced, suggesting the AIS in these cells remains functional and is potentially more excitable than wild type controls, which, when considered with the results from Purkinje neurons presented here, indicates the effects of *Tsc1* deletion on neuronal intrinsic excitability are cell-type specific.

## ACKNOWLEGMENTS

This work was supported by startup funds from the Miami University College of Arts and Sciences and by the NINDS at the National Institutes of Health: Award 1R15NS125560

## References

Andersen BB, Korbo L, Pakkenberg B (1992) A quantitative study of the human cerebellum with unbiased stereological techniques. J Comp Neurol 326:549–560.

Au KS, Williams AT, Roach ES, Batchelor L, Sparagana SP, Delgado MR, Wheless JW, Baumgartner JE, Roa BB, Wilson CM, Smith-Knuppel TK, Cheung M-YC, Whittemore VH, King TM, Northrup H (2007) Genotype/phenotype correlation in 325 individuals referred for a diagnosis of tuberous sclerosis complex in the United States. Genetics in Medicine 9:88–100.

Bailey A, Luthert P, Dean A, Harding B, Janota I, Montgomery M, Rutter M, Lantos P (1998) A clinicopathological study of autism. Brain 121 (Pt 5):889–905.

Barron T, Saifetiarova J, Bhat MA, Kim JH (2018) Myelination of Purkinje axons is critical for resilient synaptic transmission in the deep cerebellar nucleus. Sci Rep 8:1022.

Barski JJ, Dethleffsen K, Meyer M (2000) Cre recombinase expression in cerebellar Purkinje cells. Genesis 28:93–98.

Bateup HS, Johnson CA, Denefrio CL, Saulnier JL, Kornacker K, Sabatini BL (2013) Excitatory/inhibitory synaptic imbalance leads to hippocampal hyperexcitability in mouse models of Tuberous Sclerosis. Neuron 78:510.

Bean BP (2007) The action potential in mammalian central neurons. Nat Rev Neurosci 8:451–465.

Benthall KN, Ong SL, Bateup HS (2018) Corticostriatal Transmission Is Selectively Enhanced in Striatonigral Neurons with Postnatal Loss of Tsc1. Cell Rep 23:3197–3208.

Bonizzi G, Karin M (2004) The two NF-kappaB activation pathways and their role in innate and adaptive immunity. Trends Immunol 25:280–288.

Bosch MK, Carrasquillo Y, Ransdell JL, Kanakamedala A, Ornitz DM, Nerbonne JM (2015) Intracellular FGF14 (iFGF14) Is Required for Spontaneous and Evoked Firing in Cerebellar Purkinje Neurons and for Motor Coordination and Balance. J Neurosci 35:6752–6769.

Crino PB, Nathanson KL, Henske EP (2006) The tuberous sclerosis complex. N Engl J Med 355:1345–1356.

Cupolillo D, Hoxha E, Faralli A, De Luca A, Rossi F, Tempia F, Carulli D (2016) Autistic-Like Traits and Cerebellar Dysfunction in Purkinje Cell PTEN Knock-Out Mice. Neuropsychopharmacology 41:1457–1466.

Dan HC, Adli M, Baldwin AS (2007) Regulation of mammalian target of rapamycin activity in PTEN-inactive prostate cancer cells by I kappa B kinase alpha. Cancer Res 67:6263–6269.

Dan HC, Ebbs A, Pasparakis M, Van Dyke T, Basseres DS, Baldwin AS (2014) Akt-dependent activation of mTORC1 complex involves phosphorylation of mTOR (mammalian target of rapamycin) by IκB kinase α (IKKα). J Biol Chem 289:25227–25240.

de Vries PJ et al. (2023) International consensus recommendations for the identification and treatment of tuberous sclerosis complex-associated neuropsychiatric disorders (TAND). J Neurodev Disord 15:32.

Díaz-Anzaldúa A, Díaz-Martínez A (2013) [Genetic, environmental, and epigenetic contribution to the susceptibility to autism spectrum disorders]. Rev Neurol 57:556–568.

Dzhashiashvili Y, Zhang Y, Galinska J, Lam I, Grumet M, Salzer JL (2007) Nodes of Ranvier and axon initial segments are ankyrin G-dependent domains that assemble by distinct mechanisms. J Cell Biol 177:857–870.

European Chromosome 16 Tuberous Sclerosis Consortium (1993) Identification and characterization of the tuberous sclerosis gene on chromosome 16. Cell 75:1305–1315.

Fingar DC, Salama S, Tsou C, Harlow E, Blenis J (2002) Mammalian cell size is controlled by mTOR and its downstream targets S6K1 and 4EBP1/eIF4E. Genes Dev 16:1472–1487.

Fréal A, Fassier C, Le Bras B, Bullier E, De Gois S, Hazan J, Hoogenraad CC, Couraud F (2016) Cooperative Interactions between 480 kDa Ankyrin-G and EB Proteins Assemble the Axon Initial Segment. J Neurosci 36:4421–4433.

Gao Y, Gartenhaus RB, Lapidus RG, Hussain A, Zhang Y, Wang X, Dan HC (2015) Differential IKK/NF-κB Activity Is Mediated by TSC2 through mTORC1 in PTEN-Null Prostate Cancer and Tuberous Sclerosis Complex Tumor Cells. Mol Cancer Res 13:1602–1614.

Gasser A, Ho TS-Y, Cheng X, Chang K-J, Waxman SG, Rasband MN, Dib-Hajj SD (2012) An ankyrinG-binding motif is necessary and sufficient for targeting Nav1.6 sodium channels to axon initial segments and nodes of Ranvier. J Neurosci 32:7232–7243.

Gibson JM, Vazquez AH, Yamashiro K, Jakkamsetti V, Ren C, Lei K, Dentel B, Pascual JM, Tsai PT (2023) Cerebellar contribution to autism-relevant behaviors in fragile X syndrome models. Cell Rep 42:113533.

Golowasch J, Thomas G, Taylor AL, Patel A, Pineda A, Khalil C, Nadim F (2009) Membrane capacitance measurements revisited: dependence of capacitance value on measurement method in nonisopotential neurons. J Neurophysiol 102:2161–2175.

Hedstrom KL, Ogawa Y, Rasband MN (2008) AnkyrinG is required for maintenance of the axon initial segment and neuronal polarity. J Cell Biol 183:635–640.

Ho TS-Y, Zollinger DR, Chang K-J, Xu M, Cooper EC, Stankewich MC, Bennett V, Rasband MN (2014) A hierarchy of ankyrin-spectrin complexes clusters sodium channels at nodes of Ranvier. Nat Neurosci 17:1664–1672.

Holmes GL, Stafstrom CE, Tuberous Sclerosis Study Group (2007) Tuberous sclerosis complex and epilepsy: recent developments and future challenges. Epilepsia 48:617–630.

Inoki K, Guan K-L (2009) Tuberous sclerosis complex, implication from a rare genetic disease to common cancer treatment. Hum Mol Genet 18:R94–100.

Ito M, Yoshida M, Obata K (1964) Monosynaptic inhibition of the intracerebellar nuclei induced from the cerebellar cortex. Experientia 20:575–576.

Jenkins SM, Bennett V (2001) Ankyrin-G coordinates assembly of the spectrin-based membrane skeleton, voltage-gated sodium channels, and L1 CAMs at Purkinje neuron initial segments. J Cell Biol 155:739–746.

Jeste SS, Sahin M, Bolton P, Ploubidis GB, Humphrey A (2008) Characterization of autism in young children with tuberous sclerosis complex. J Child Neurol 23:520–525.

Kalume F, Yu FH, Westenbroek RE, Scheuer T, Catterall WA (2007) Reduced sodium current in Purkinje neurons from Nav1.1 mutant mice: implications for ataxia in severe myoclonic epilepsy in infancy. J Neurosci 27:11065–11074.

Khaliq ZM, Raman IM (2006) Relative Contributions of Axonal and Somatic Na Channels to Action Potential Initiation in Cerebellar Purkinje Neurons. J Neurosci 26:1935–1944.

König H-G, Schwamborn R, Andresen S, Kinsella S, Watters O, Fenner B, Prehn JHM (2017) NF-κB regulates neuronal ankyrin-G via a negative feedback loop. Sci Rep 7:42006.

Kwiatkowski DJ (2003) Rhebbing up mTOR: new insights on TSC1 and TSC2, and the pathogenesis of tuberous sclerosis. Cancer Biol Ther 2:471–476.

Kwiatkowski DJ, Zhang H, Bandura JL, Heiberger KM, Glogauer M, el-Hashemite N, Onda H (2002) A mouse model of TSC1 reveals sex-dependent lethality from liver hemangiomas, and up-regulation of p70S6 kinase activity in Tsc1 null cells. Human Molecular Genetics 11:525–534.

Lawson RJ, Lipovsek NJ, Brown SP, Jena AK, Osko JJ, Ransdell JL (2024) Selective deletion of Tsc1 from mouse cerebellar Purkinje neurons drives sex-specific behavioral impairments linked to autism. Front Behav Neurosci 18 Available at: https://www.frontiersin.org/journals/behavioral-neuroscience/articles/10.3389/fnbeh.2024.1474066/full [Accessed December 23, 2024].

Levin SI, Khaliq ZM, Aman TK, Grieco TM, Kearney JA, Raman IM, Meisler MH (2006) Impaired motor function in mice with cell-specific knockout of sodium channel Scn8a (NaV1.6) in cerebellar purkinje neurons and granule cells. J Neurophysiol 96:785–793.

Liu F, Xia Y, Parker AS, Verma IM (2012) IKK biology. Immunol Rev 246:239–253.

Madisen L, Zwingman TA, Sunkin SM, Oh SW, Zariwala HA, Gu H, Ng LL, Palmiter RD, Hawrylycz MJ, Jones AR, Lein ES, Zeng H (2010) A robust and high-throughput Cre reporting and characterization system for the whole mouse brain. Nat Neurosci 13:133–140.

Marcotte L, Crino PB (2006) The neurobiology of the tuberous sclerosis complex. Neuromolecular Med 8:531–546.

Meeks JP, Mennerick S (2007) Action potential initiation and propagation in CA3 pyramidal axons. J Neurophysiol 97:3460–3472.

Milescu LS, Bean BP, Smith JC (2010) Isolation of somatic Na+ currents by selective inactivation of axonal channels with a voltage prepulse. J Neurosci 30:7740–7748.

Ng S, Wu Y-T, Chen B, Zhou J, Shen H-M (2011) Impaired autophagy due to constitutive mTOR activation sensitizes TSC2-null cells to cell death under stress. Autophagy 7:1173–1186.

Northrup H (1992) Tuberous sclerosis complex: genetic aspects. J Dermatol 19:914–919.

Obata K, Ito M, Ochi R, Sato N (1967) Pharmacological properties of the postsynaptic inhibition by Purkinje cell axons and the action of γ-aminobutyric acid on Deiters neurones. Exp Brain Res 4:43–57.

Obata K, Takeda K, Shtnozaki H (1970) Further study on pharmacological properties of the cerebellar-induced inhibition of Deiters neurones. Exp Brain Res 11:327–342.

Orban PC, Chui D, Marth JD (1992) Tissue- and site-specific DNA recombination in transgenic mice. Proc Natl Acad Sci U S A 89:6861–6865.

Palkovits M, Magyar P, Szentágothai J (1972) Quantitative histological analysis of the cerebellar cortex in the cat. IV. Mossy fiber-Purkinje cell numerical transfer. Brain Res 45:15–29.

Peter S, Ten Brinke MM, Stedehouder J, Reinelt CM, Wu B, Zhou H, Zhou K, Boele H-J, Kushner SA, Lee MG, Schmeisser MJ, Boeckers TM, Schonewille M, Hoebeek FE, De Zeeuw CI (2016) Dysfunctional cerebellar Purkinje cells contribute to autism-like behaviour in Shank2-deficient mice. Nat Commun 7:12627.

Port RG, Gandal MJ, Roberts TPL, Siegel SJ, Carlson GC (2014) Convergence of circuit dysfunction in ASD: a common bridge between diverse genetic and environmental risk factors and common clinical electrophysiology. Front Cell Neurosci 8:414.

Ransdell JL, Brown SP, Xiao M, Ornitz DM, Nerbonne JM (2024) In Vivo Expression of an SCA27A-linked FGF14 Mutation Results in Haploinsufficiency and Impaired Firing of Cerebellar Purkinje Neurons. bioRxiv:2024.10.25.620253.

Ransdell JL, Dranoff E, Lau B, Lo W-L, Donermeyer DL, Allen PM, Nerbonne JM (2017) Loss of Navβ4-Mediated Regulation of Sodium Currents in Adult Purkinje Neurons Disrupts Firing and Impairs Motor Coordination and Balance. Cell Rep 19:532–544.

Saito H, Tsumura H, Otake S, Nishida A, Furukawa T, Suzuki N (2005) L7/Pcp-2-specific expression of Cre recombinase using knock-in approach. Biochem Biophys Res Commun 331:1216–1221.

Springer SJ, Burkett BJ, Schrader LA (2014) Modulation of BK channels contributes to activity-dependent increase of excitability through MTORC1 activity in CA1 pyramidal cells of mouse hippocampus. Front Cell Neurosci 8:451.

Stoodley CJ (2014) Distinct regions of the cerebellum show gray matter decreases in autism, ADHD, and developmental dyslexia. Front Syst Neurosci 8:92.

Stoodley CJ, D’Mello AM, Ellegood J, Jakkamsetti V, Liu P, Nebel MB, Gibson JM, Kelly E, Meng F, Cano CA, Pascual JM, Mostofsky SH, Lerch JP, Tsai PT (2017) Altered cerebellar connectivity in autism spectrum disorders and rescue of autism-related behaviors in mice. Nat Neurosci 20:1744–1751.

Stuart G, Häusser M (1994) Initiation and spread of sodium action potentials in cerebellar Purkinje cells. Neuron 13:703–712.

Tee AR, Fingar DC, Manning BD, Kwiatkowski DJ, Cantley LC, Blenis J (2002) Tuberous sclerosis complex-1 and -2 gene products function together to inhibit mammalian target of rapamycin (mTOR)-mediated downstream signaling. Proc Natl Acad Sci U S A 99:13571–13576.

Tee AR, Manning BD, Roux PP, Cantley LC, Blenis J (2003) Tuberous sclerosis complex gene products, Tuberin and Hamartin, control mTOR signaling by acting as a GTPase-activating protein complex toward Rheb. Curr Biol 13:1259–1268.

Tsai PT, Hull C, Chu Y, Greene-Colozzi E, Sadowski AR, Leech JM, Steinberg J, Crawley JN, Regehr WG, Sahin M (2012) Autistic-like behaviour and cerebellar dysfunction in Purkinje cell Tsc1 mutant mice. Nature 488:647–651.

Tsai PT, Rudolph S, Guo C, Ellegood J, Gibson JM, Schaeffer SM, Mogavero J, Lerch JP, Regehr W, Sahin M (2018) Sensitive Periods for Cerebellar-Mediated Autistic-like Behaviors. Cell Rep 25:357–367.e4.

Wang SS-H, Kloth AD, Badura A (2014) The cerebellum, sensitive periods, and autism. Neuron 83:518–532.

Witter L, Rudolph S, Pressler RT, Lahlaf SI, Regehr WG (2016) Purkinje Cell Collaterals Enable Output Signals from the Cerebellar Cortex to Feed Back to Purkinje Cells and Interneurons. Neuron 91:312–319.

Wiznitzer M (2004) Autism and tuberous sclerosis. J Child Neurol 19:675–679.

Wullschleger S, Loewith R, Hall MN (2006) TOR signaling in growth and metabolism. Cell 124:471–484.

Xiao M, Bosch MK, Nerbonne JM, Ornitz DM (2013) FGF14 localization and organization of the axon initial segment. Mol Cell Neurosci 56:393–403.

Yang Y, Ogawa Y, Hedstrom KL, Rasband MN (2007) betaIV spectrin is recruited to axon initial segments and nodes of Ranvier by ankyrinG. J Cell Biol 176:509–519.

Yoon S, Piguel NH, Penzes P (2022) Roles and mechanisms of ankyrin-G in neuropsychiatric disorders. Exp Mol Med 54:867–877.

Zhang X-M, Ng AH-L, Tanner JA, Wu W-T, Copeland NG, Jenkins NA, Huang J-D (2004) Highly restricted expression of Cre recombinase in cerebellar Purkinje cells. Genesis 40:45–51.

Zhou D, Lambert S, Malen PL, Carpenter S, Boland LM, Bennett V (1998) AnkyrinG is required for clustering of voltage-gated Na channels at axon initial segments and for normal action potential firing. J Cell Biol 143:1295–1304.

