## Supplemental Table and Figures for "*Tsc1* Deletion in Purkinje Neurons Disrupts the Axon Initial Segment, Impairing Excitability and Cerebellar Function"

### *Supplemental Table 1 Brown et al.*

| **Current-clamp (slice)** | **Controls** | ***Tsc1*$^{mut/mut}$** | ***Tsc1*$^{mut/+}$** |
|---|---|---|---|
| Animal Numbers (male ; female) | 6 ; 7 | 3 ; 3 | 2 ; 4 |
| Age Range (days old) | 38-60 | 38-46 | 40-53 |
| **Immunofluoresence** | **Controls** | ***Tsc1*$^{mut/mut}$** | ***Tsc1*$^{mut/+}$** |
| Animal Numbers (male ; female) | 3 ; 1 | 1 ; 3 | 2 ; 1 |
| Age Range (days old) | 35-78 | 42-52 | 42-73 |
| **Voltage-clamp (slice activation recordings)** | **Controls** | ***Tsc1*$^{mut/mut}$** | - |
| Animal Numbers (male ; female) | 4 ; 2 | 5 ; 1 | - |
| Age Range (days old) | 43-53 | 46-50 | - |

*Supplemental Figure 1 Brown et al.*

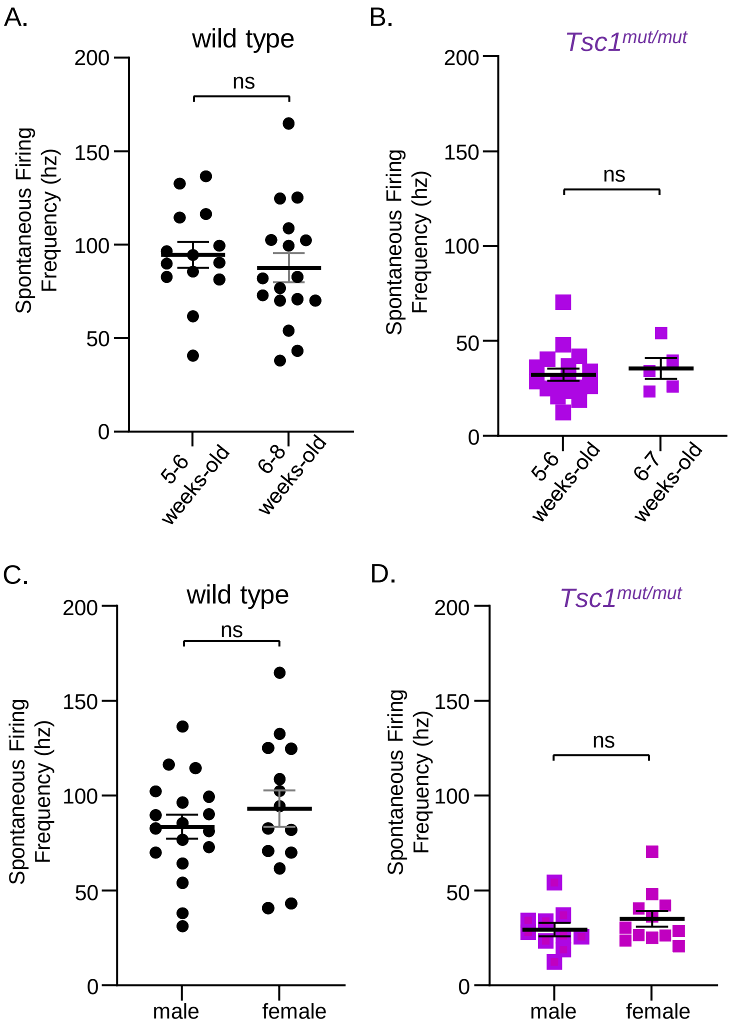


**In wild type and *Tsc1*$^{mut/mut}$ Purkinje neurons, mean firing frequency is not different across age and sex.**

**A. B.** Current-clamp experiments were performed on wild type (**A.**) and*Tsc1*$^{mut/mut}$ (**B.**) Purkinje neurons isolated from younger (≤6 week-old) and older (>6 weeks-old) animals. Within each genotype, mean (± SEM) firing frequency was similar between these age groups (unpaired Student's t-test). Within the wild type (**C.**) and *Tsc1*$^{mut/mut}$ (**D.**) genotypes, mean (± SEM) firing frequency was also found to be similar between male and female animals (unpaired Student's t-test).

*Supplemental Figure 2 Brown et al.*

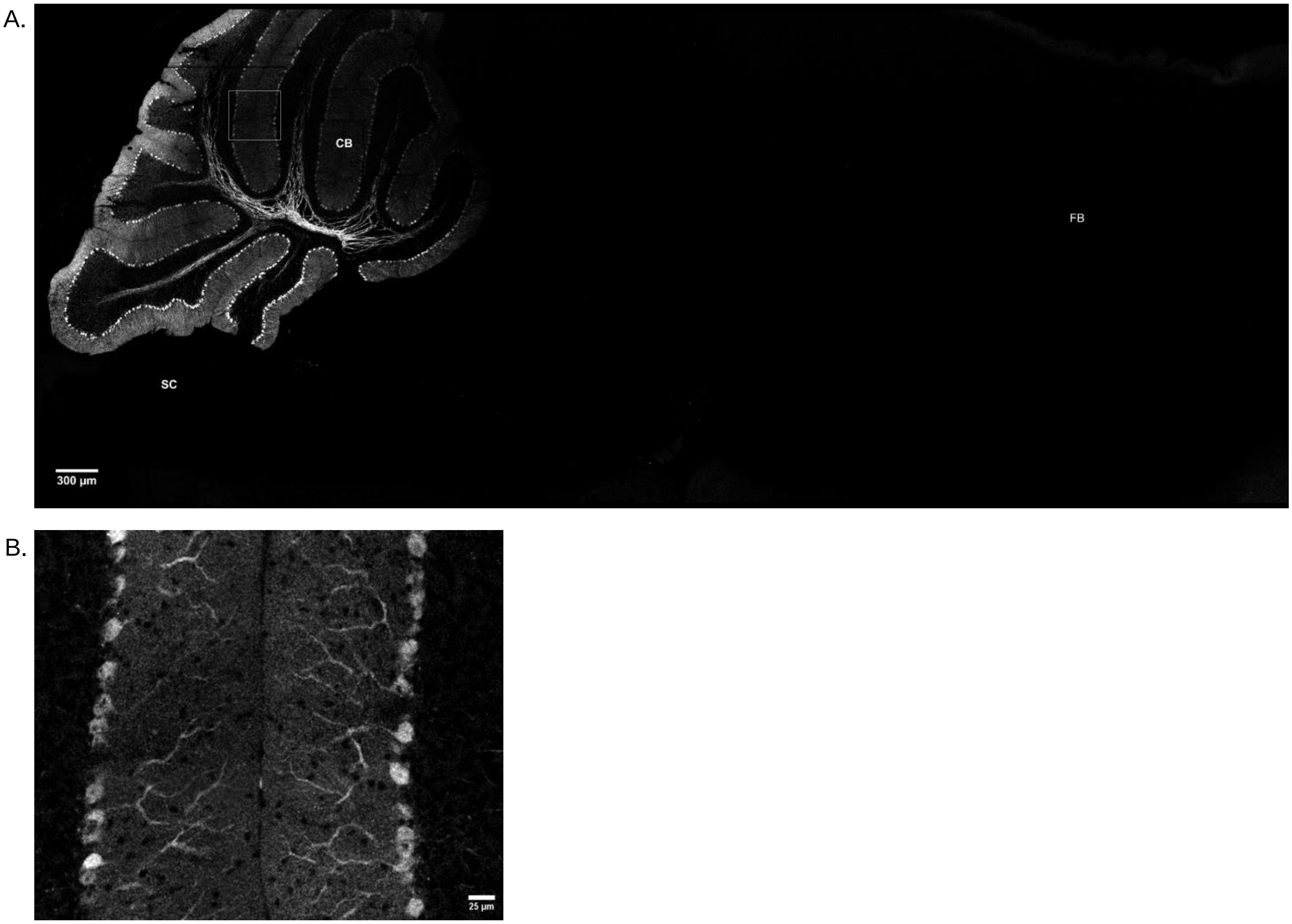


**L7/Pcp2 Cre-mediated tdTomato fluorescence is isolated to the somata and neurites of cerebellar Purkinje neurons.**

**A**. Fluorescent micrograph of a sagittal brain section from an adult L7/Pcp2 Cre-positive; Ai14 Cre-reporter mouse revealing localized Cre-mediated tdTomato fluorescence in the cerebellum and an absence of Cre-mediated fluorescence in the spinal cord (SC) and forebrain (FB). Image acquired using a 20X objective. Panel **B.** presents an enhanced view of the cerebellar Purkinje neuron and molecular layers of sections of lobules V and VI marked in panel A. using a white dashed box.

*Supplemental Figure 3 Brown et al.*

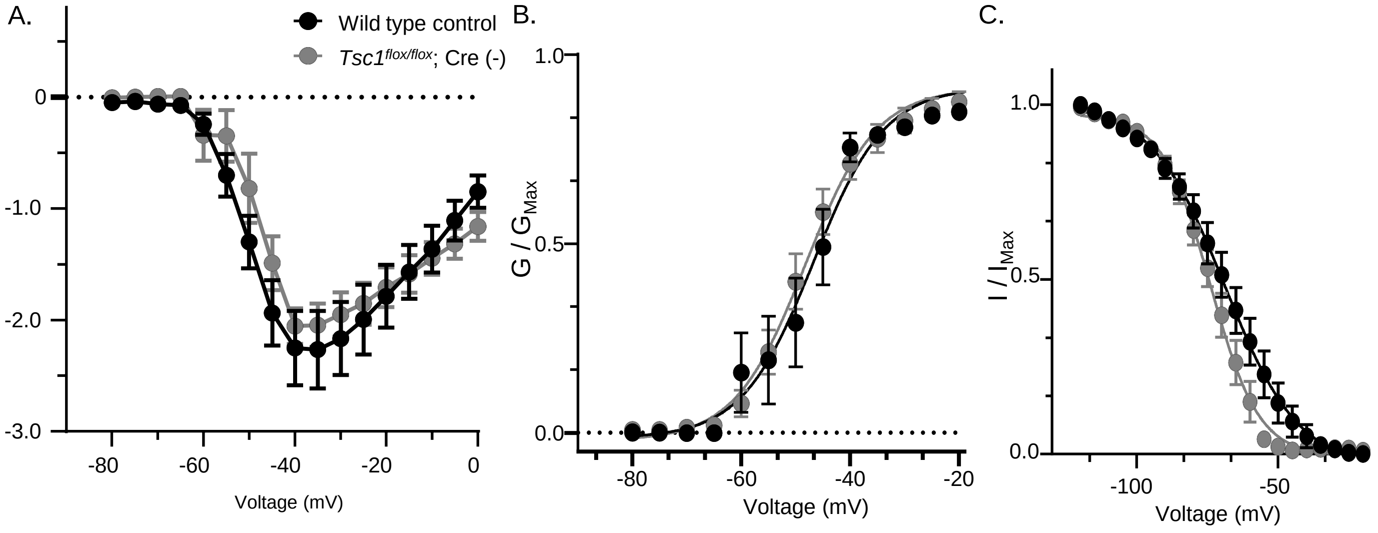


**In adult Purkinje neurons, Nav currents properties are similar across control groups**

**A.** The mean (± SEM) peak $I_{NaT}$ values are plotted against activating voltage steps (see Figure 4B) for wild type and *Tsc1$^{flox/flox}$*; Cre (-) control groups, revealing no difference in the amplitude of evoked $I_{NaT}$. **B.** $I_{NaT}$ values were used to calculate Nav conductance and the mean (± SEM) normalized conductance values were plotted against activating voltages steps revealing no differences in the voltage-dependence of Nav conductance activation. Data were fitted using a1$^{st}$ order Boltzmann sigmoidal equation. $V_{1/2}$ values from Boltzmann sigmoidal fits equal -46.5 mV and -48.1 mV for wild type and *Tsc1$^{flox/flox}$*;Cre (-) control groups, respectively. **C.** To assess the voltage-dependence of Nav current steady-state inactivation, mean (± SEM) normalized peak $I_{NaT}$ values, evoked during a common depolarizing voltage step to -20 mV, are plotted against the prior conditioning voltage (see Methods). These plots reveal the voltage-dependence of Nav current steady-state inactivation is similar between control groups in adult Purkinje neurons. $V_{1/2}$ values from Boltzmann sigmoidal fits equal -69.1 mV and -73.9 mV for wild type and *Tsc1$^{flox/flox}$*;Cre (-) control groups, respectively.

*Supplemental Figure 4 Brown et al.*

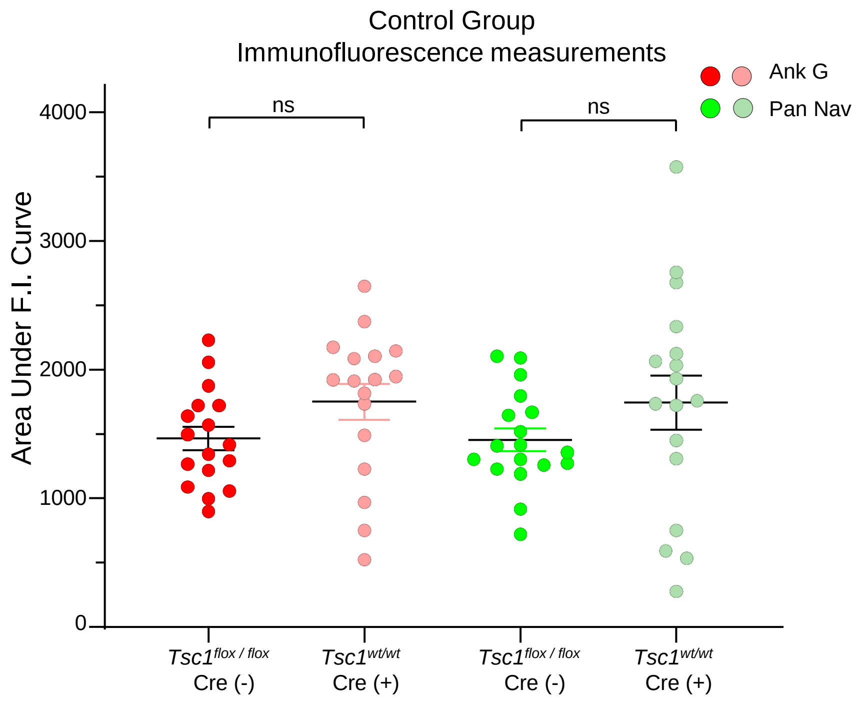


**Immunofluorescence integrated intensity measures are not different between control groups.**

Fluorescence intensity values reflecting anti-ankyrinG (*red*) and anti-pan Nav (*green*) channel immunofluorescence were measured from line scans drawn along Purkinje cell axon initial segments. These values were plotted against distance from the cell somata. For each cell's fluorescence intensity plot, the area under the curve (AUC) was measured and compared across control groups. Unpaired Student's t-tests reveal AUC values are not significantly different across control groups for anti-ankyrinG and anti-pan Nav channel intensity plots.
